# Interpreting machine learning models to investigate circadian regulation and facilitate exploration of clock function

**DOI:** 10.1101/2021.02.04.429826

**Authors:** Laura-Jayne Gardiner, Rachel Rusholme-Pilcher, Josh Colmer, Hannah Rees, Juan Manuel Crescente, Anna Paola Carrieri, Susan Duncan, Edward O. Pyzer-Knapp, Ritesh Krishna, Anthony Hall

## Abstract

The circadian clock is an important adaptation to life on earth. Here, we use machine learning to predict complex temporal circadian gene expression patterns in *Arabidopsis*. Most significantly, we classify circadian genes using DNA sequence features generated from public genomic resources, with no experimental work or prior knowledge needed. We use model explanation to rank DNA sequence features, observing transcript-specific combinations of potential circadian regulatory elements that discriminate temporal phase of expression. Model interpretation/explanation provides the backbone of our methodological advances, giving insight into biological processes and experimental design. Next, we use model interpretation to optimize sampling strategies when we predict circadian transcripts using reduced numbers of transcriptomic timepoints, saving both time and money. Finally, we predict the circadian time from a single transcriptomic timepoint, deriving novel marker transcripts that are most impactful for accurate prediction, this could facilitate the identification of altered clock function from existing datasets.

## Introduction

The circadian clock is an internal molecular 24-hour timer that is a critical adaptation to life on Earth. It temporally orchestrates physiology, biochemistry and metabolism across the day/night cycle. As a result, it regulates many traits associated with fitness and survival [1,2]. The clock is a well characterised transcriptional regulatory network which drives complex, widespread and robust patterns of temporal gene expression [3,4]. However, our understanding of such complex transcriptional regulatory systems is limited by our ability to assay them, requiring the generation of long high-resolution time-series datasets.

In plants, much of our understanding of circadian regulation, comes from our study of the model plant *Arabidopsis thaliana.* This has yielded a plethora of public multi-omic resources [5,6,7] that can be re-analysed to give new insights into the roles and functions of complex regulatory networks. In this study, we use newly generated datasets, published temporal datasets [8,9,10] (Table S1) and *Arabidopsis* genomes, in combination with machine learning (ML) approaches (see Glossary for definitions of terms), to make predictions about circadian gene regulation and expression patterns. Critically, we advance existing approaches using explainable AI algorithms and interpretation of our models (Glossary), such methods help us to understand the predictions made by ML models. In this case, giving insight into biological processes and experimental design alongside our predictions. Clarity with respect to how a model makes its predictions, we propose, will also generate confidence and trust in the model, promoting its usage. We use the *Arabidopsis* circadian clock as an example of a complex transcriptional regulatory network since some of its key regulatory elements are already known, allowing validation of our findings with experimental evidence.

Circadian gene expression rhythms reflect a variety of waveform shapes with a characteristic periodicity of ~24h [11]. Recent computational methods for identifying these rhythms from transcriptomic time course datasets have achieved circadian gene classification with as few as 3-6 timepoints (saving time for sampling and money for sequencing) [12]. However, some of the most popular approaches describe optimal sampling strategies for the identification of rhythms running with >3 days of data and 2-hourly sampling resolution [13, 14]. We propose that this is partly due to concern for the loss of information as a result of down sampling. Since the cost implications of this are high, our focus is on designing trusted down-sampling strategies for capturing circadian oscillations using a non-optimal number of timepoints. As such, firstly, we develop ML models that not only classify circadian expression patterns using iteratively lower numbers of transcriptomic timepoints improving accuracy compared to the state-of-the-art. But moreover, we use model interpretation to quantify the best transcriptomic timepoints for sampling. We believe that this predictive insight on when to sample will be a valuable reference for experimental biologists when planning experiments.

Next, we re-define the field, developing ML models that distinguish circadian transcripts using no transcriptomic timepoint information, and instead using only DNA sequence features (Glossary). The theory supporting this is that a major mechanism of (circadian or otherwise) gene expression control is through transcription factor binding to regulatory DNA sequence. Considering previous work in *Arabidopsis* it is likely that the promoter, 5’UTR and the first part of the coding region are the most useful locations for transcription factor binding site (TFBS) detection [15]. Genes expressed with similar patterns are more likely to be controlled by similar sets of TFBSs. In addition, small RNAs (sRNAs), comprising microRNAs (miRNAs) and small interfering RNAs (siRNAs) are thought to affect transcript abundance via post-transcriptional regulation of mRNA [16]. Plant miRNAs predominately bind to the coding regions of mRNA, and to a lesser extent 5′UTR and 3′UTR regions [17,18]. As such, we consider both coding and non-coding regions to classify circadian genes using DNA sequence. Our DNA sequence features are profiles of *k-*mer-based motif representations that are identified *de novo* and embody a comprehensive picture of TFBS, sRNA/RNA binding sites and other sequence-based regulatory elements, since we incorporate the promoter, 5’UTR, 3’UTR and coding regions.

A key strength of our DNA-sequence based approach is that we classify circadian transcripts using *k-* mer-based motif representations generated from pre-existing public genomic resources with no experimental work or prior knowledge of regulatory elements needed. Computational regulatory motif discovery methods typically search for overrepresented words across DNA sequences using methods such as Expectation Maximization (EM) and Gibbs sampling [19,20,21,22]. Approaches are typically limited by a requirement for input information e.g., co-expressed genes, site abundance, number of sites per sequence or a fixed motif length [23,24,25]. Furthermore, Artificial Intelligence (AI) has been used to predict transcriptomic profiles directly using features such as DNA sequence or epigenetic marks. These features typically include representations of TFBS [26,27], enhancers [28], histone modifications [29] or open chromatin regions [30]. However, again, these approaches typically require experimental data or prior knowledge of regulatory elements that our approach does not need, or they focus on single gene expression states and do not consider complex patterns, as our methods do.

Additionally, AI-based work in the field of expression prediction has largely lacked comprehensive model explanation [31]. Here, we expose the potential, alongside our DNA-sequence based predictive model, to use explainable AI to discover regulatory motifs and explore their functional consequences. We exploit model explanation to identify, on a transcript-by-transcript basis, the ranked regulatory sequences that guide the classification of its expression pattern as circadian. We identify both small and larger combinations of regulatory elements that, in combination, give a larger overall impact on gene classification. These regulatory sequences are candidate causal genetic features that could control gene expression and allow us to understand the regulatory mechanisms governing circadian expression patterns and even manipulate its regulation, focused here on circadian rhythmicity. Ultimately, we use model explanation to generate and validate hypotheses *in silico*, facilitating both gene expression prediction and derivative regulatory element discovery.

Finally, assaying circadian clock function, as opposed to simply identifying transcript rhythmicity, has been a major challenge for the study of the circadian regulation in organisms ranging from mammals to plants. Recent work applied ML to circadian time course transcriptomic datasets from human blood, to predict the phase of the endogenous circadian clock (circadian time, CT), using a single time point from a set of marker genes [32,33]. This allows the use of one time point to identify altered clock function e.g., due to disease or environmental conditions. An equivalent major challenge exists in plant sciences. As such, we use ML to predict the circadian time in *Arabidopsis* from a single transcriptomic timepoint using marker genes. To advance previous offerings, we identify novel marker genes as part of our interpretable approach ensuring that they represent a diverse range of temporal patterns with consistent amplitudes across datasets to facilitate accurate and robust phase prediction irrespective of sample phase. Counter-intuitively our marker genes do not include the core clock genes used in previous studies for time prediction [34]. Taken together, these tools constitute a suite of informative resources for both experimental biologists and the interpretation of further circadian datasets.

## Results and Discussion

### ML model interpretation optimizes timepoint down-sampling to define circadian transcripts

We used MetaCycle as our baseline for detecting circadian signals in dense time-series transcriptomic data [13]. MetaCycle is one of the most well-maintained and accessible tools within the community incorporating a variety of the most widely used methods ARSER [35], JTK_CYCLE [36] and Lomb-Scargle [37] and integrating their results so that rhythmic prediction is a cumulation of different statistical approaches. We ran MetaCycle (see Methods) on a published *Arabidopsis* time-series transcriptomic dataset generated by [8], which was sampled every 4-hours for 48-hours, starting 24-hours after transfer to constant conditions (LL) (Table S1). The data was processed to produce normalized counts per transcript (see Methods). MetaCycle classified 9,394 out of 44,963 transcripts as circadian (q<0.05), with 7,734 denoted as high confidence (q<0.02) (Supplementary Note 1). We trained a series of ML classifiers to predict if a transcript was circadian or non-circadian in a binary classification system using 7,734 of the least likely candidates to be circadian (q>0.99) labelled by MetaCycle alongside the 7,734 highly circadian transcripts (q<0.02) (see Methods; Glossary; Supplementary Note 2). For the ML models we report the F1 scores that measure the accuracy of the model on a scale of 0 to 1, with 1 being most accurate (Glossary). Considering all 12 transcriptomic time points, the best model was generated with LightGBM after optimization (Methods; Figure S1a, Table S2) with: an F1 score of 0.999 on the training data, an F1 score of 0.955 on the (held out) test data and a mean F1 cross validation score of 0.939 (Glossary). Our confusion matrix (Figure S1b; Glossary) highlights consistently high accuracy of our model irrespective of the class that is being predicted (circadian/non-circadian).

Our best ML model (LightGBM) was able to assign a matching circadian/non-circadian label to the majority of the transcripts that MetaCycle labelled. Overall, there is good agreement between our model and MetaCycle. However, the overlap was not 100% so we examined the small proportion of transcripts that were “inaccurately” classified. We found that the “inaccurately” classified cases by our ML model were more likely to be intermediate or border-line cases for MetaCycle (Figure 1) or edge cases e.g., with slightly longer period lengths (Figure S1). We deduced this because cases rejected by MetaCycle as circadian but accepted by the ML (false positives-FP) had significantly lower (MetaCycle derived) p-values than the cases that were rejected by both MetaCycle and ML (true negatives-TN) (p<0.0001, t=6.8795, df=7753). Conversely, cases accepted by MetaCycle as rhythmic but rejected by ML (false negatives-FN) had higher (MetaCycle derived) p-values than cases categorised as rhythmic by both MetaCycle and ML (true positives-TP) (p<0.0001, t=5.7744, df=7711) (Figure 1a). Additionally, cases rejected by MetaCycle as circadian but accepted by the ML (FP) have significantly lower relative amplitudes compared to the TP calls where both methods agree (p<0.0001, t=8.3845, df=7732). Conversely, cases accepted by Metacycle as rhythmic but rejected by ML (FN), had a significantly higher relative amplitude than the true negative calls (p=0.036, t=2.0924, df=7732) (Figure 1b). This also highlights that the ML model is not simply using high and low expression levels to discriminate circadian and non-circadian status of transcripts.

**Figure 1.**
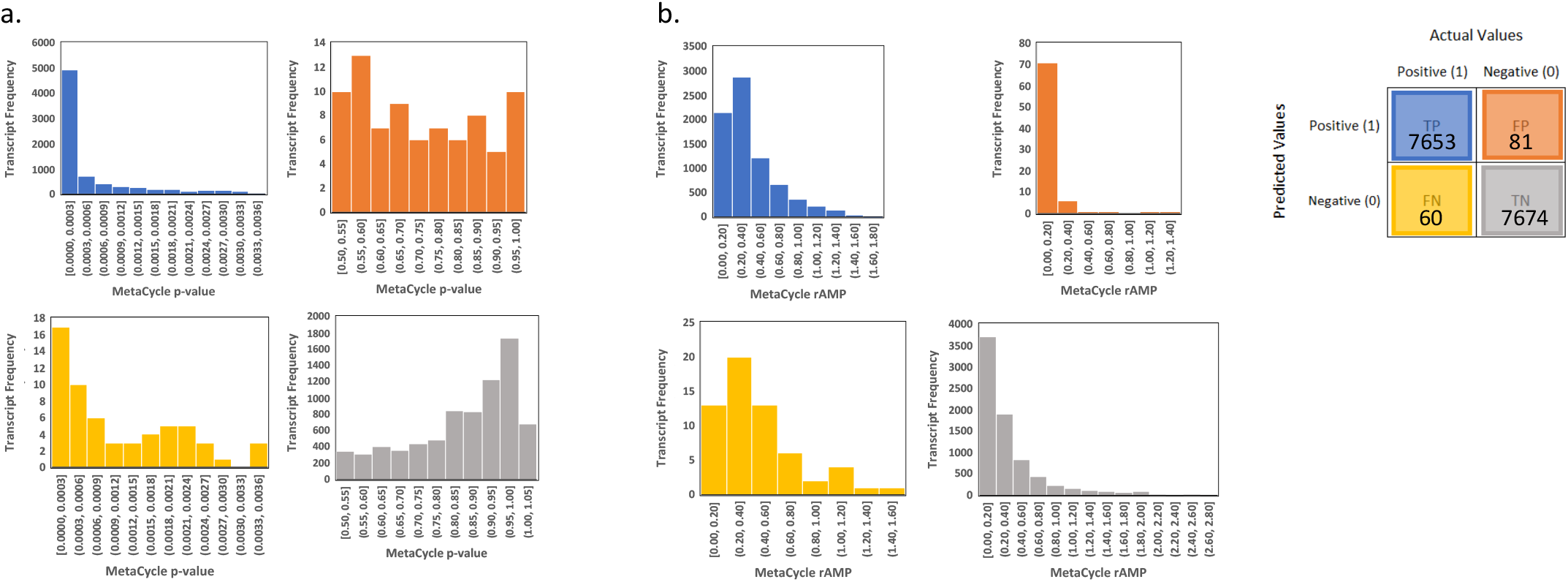
Arabidopsis circadian/non-circadian comparative ML binary classification analysis with 12 transcriptomic timepoints. Class 0=Non-circadian and Class 1=Circadian. Histograms in **(a-b)** all relate to the best model from Figure S1a that was generated using LightGBM, the histograms are colour coded as per the confusion matrix shown in the legend to the right i.e. showing where our model assigned True Positive labels (TP), False Positive labels (FP), False Negative labels (FN) and True Negative labels (TN). The histograms show the frequency of transcripts that had various **(a)** p-values or **(b)** relative amplitudes assigned to them by Metacycle.

We assessed the effect of reducing the number of transcriptomic timepoints on the accuracy of our classification of circadian/non-circadian transcripts. For our best ML model (derived using 12 timepoints), we reduced the number of timepoints (or features) sequentially from 12 down to 3. To obtain each of the interim reduced sets of timepoints from 12 to 3, we used well-known feature selection tools chi-square and eli5 (Glossary) and compared these against testing every possible feature combination for the timepoint number (see Methods). The method of trialling every possible feature combination for each reduced timepoint number enabled us to most accurately classify transcripts as circadian/non-circadian (Figure 2a). Using this approach with 6 timepoints, we achieved a mean classification F1 accuracy score of 0.886 on cross validation and a score of 0.792 using only 3 time points (Table S3). Table S3 also highlights, for these most accurate models, that we have consistently high accuracy irrespective of the class that is being predicted (circadian/non-circadian). Using model interpretation i.e., identifying the combinations of features that gave the highest accuracies, we were able to define the most optimal sampling strategies for the different numbers of timepoints. For selection of 6 or more timepoints, the best combinations tended to be consecutive timepoints extending across the intersect of day 1 and day 2. In contrast, when selecting low numbers of timepoints, more accurate classifications were made when timepoints were spaced across a single day (Figure 2b). Figure 2c highlights this showing the best combination of reduced timepoints in each category 12-3 for the example transcript phytochrome A (*PHYA*).

**Figure 2.**
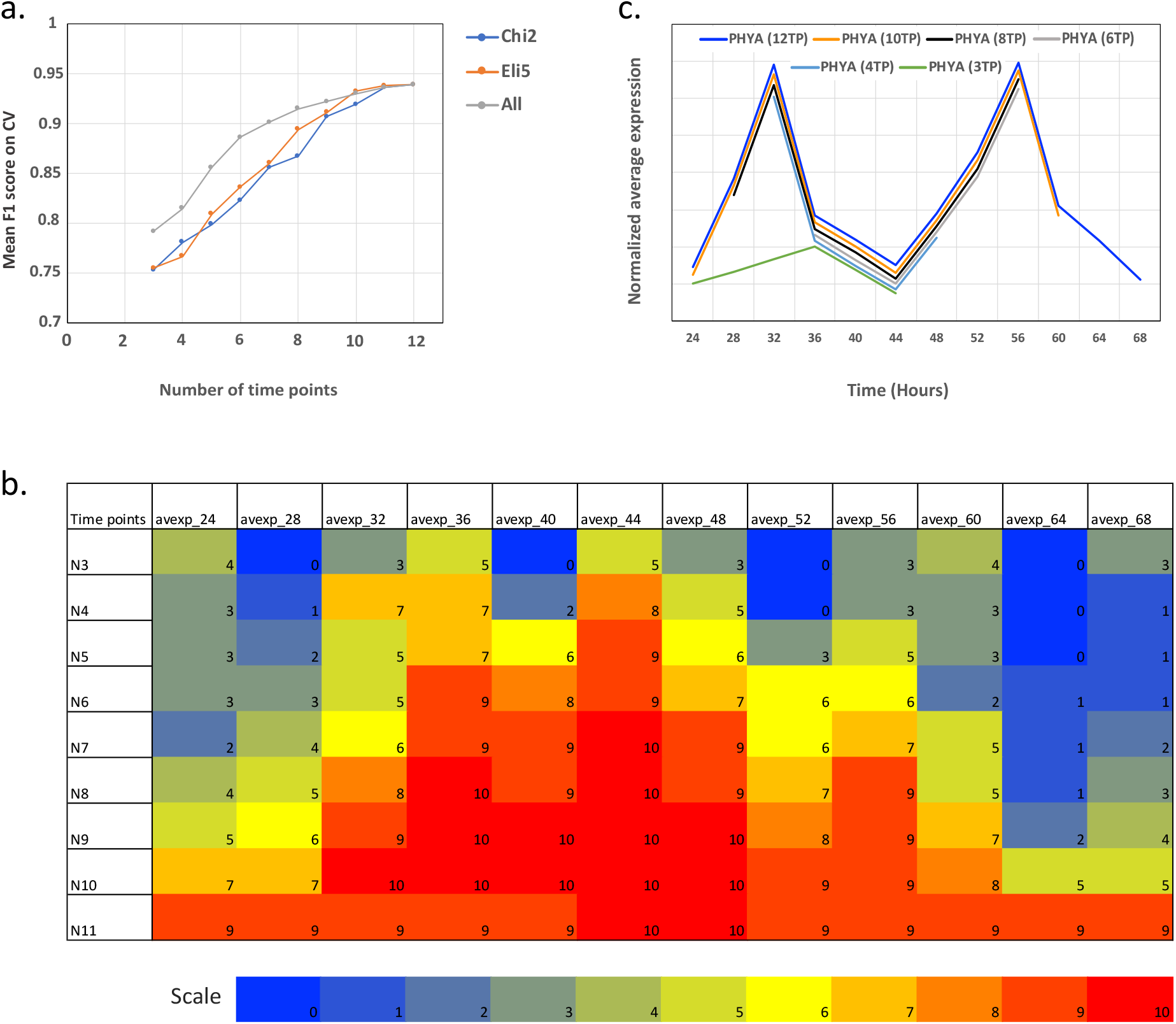
Arabidopsis circadian/non-circadian comparative ML binary classification analysis to reduce the number of transcriptomic timepoints. For our best ML model, we reduced the number of timepoints sequentially from 12 down to 3. **(a)** To obtain each reduced set of timepoints, we compare using chi-square (Chi2) and eli5 (Eli5) feature selection with the best set comparing every possible random feature combination (All). Here we show the best F1 score after 5-fold cross validation for each set of reduced timepoints. **(b)** Detailing the 10 best combinations of features that gave the highest accuracy or F1 score for each reduced set of timepoints. Labels N3-N11 show the number of reduced timepoints. Labels avexp_24-avexp_68 show the sampling times. Counts 0-10 represent the number of times each timepoint appeared in the 10 best combinations of features. **(c)** For the example gene *PHYA* showing a line plot of the gene’s expression values across the best combination of reduced timepoints in each category 12-3. Expression values are uniformly reduced by ~5% for each reduced timepoint combination to allow separation of lines for visualization.

In order to test how generalizable our model is on unseen data (Glossary), we used the most accurate model for the reduced set of 3 timepoints (timepoints 36, 48 and 60) for the binary classification of, firstly, a second *Arabidopsis* transcriptomic time-series dataset developed by [9] and secondly, a newly developed wheat transcriptomic dataset representing a divergent plant species from *Arabidopsis* (Table S1). These additional unrelated test datasets represent different sampling strategies and experimental setups (see Methods). Both test datasets were processed bioinformatically as per our original [8] dataset (see Methods). For the *Arabidopsis* [9] dataset, the timepoints did not match those used to train our model; sampling started 2 hours after exposure to constant light (rather than 24 hours after) and samples were taken every 3 hours instead of every 4. As such, we selected the closest times to those that were used to train our model according to time of day relative to dawn (timepoints 11, 23 and 35). Even so, the F1 score (representing accuracy) for classification of this gene set was relatively high at 0.714, amounting to a decrease in accuracy of only 0.08 compared to the dataset that the model was trained on. For the wheat dataset, sampling started 24 hours after exposure to constant light and measurements were taken every 2 hours instead of every 4. Therefore, here, matching the time of day relative to dawn, we were able to select equivalent timepoints (12, 24 and 36 hours) and the F1 score was slightly higher at 0.769 amounting to a decrease of only 0.02 on a highly divergent species. The model therefore generalizes well irrespective of the sample’s species, particularly with matched timepoints relative to dawn.

We compared our timepoint reduction analysis using ML to a range of analyses representing the state-of-the-art across the different timepoint numbers. MetaCycle requires a minimum of 6 timepoints for circadian analysis, and benefits from these timepoints being evenly sampled across the chosen time period [13]. As such, we reduced timepoints from 12 to 6 to enable comparison including evenly spaced sampling patterns; 4hourly/1day, 8hourly/2days versus the best suggested sampling times from our ML analysis (4hourly/1day from 36-56 hours from Figure 2b and 2c). The reduction to 6 timepoints significantly decreased the number of positive circadian gene calls by MetaCycle that were conserved with the 12 timepoint analysis, independently of the sampling technique used. In fact, the highest proportion of the 9,394 circadian genes identified with 12 timepoints by MetaCycle that were also identified with 6 timepoints (p<0.05) was 63.7% (Table S4). This accuracy is still ~25% lower than the F1 score we achieved with 6 timepoints and our best ML model (Table S3). Furthermore, when comparing the F1 score of our 3-timepoint ML model it was more appropriate to use a 3-timepoint state-of-the-art analysis performed by Spörl et al. [12]. Table S4 highlights that we achieve a 12% higher accuracy with only 3 timepoints in a like-for-like comparison with Spörl et al. [12]. This accuracy improvement is in addition to the experimental design insight that we provide.

### Circadian genes can be classified using *de novo* generated DNA sequence-based *k*-mer spectra

We investigated if it was possible to eliminate transcriptomic timepoints completely and use DNA sequence features alone to classify transcripts as circadian/non-circadian. To achieve this, we generated *k*-mer profiles *de-novo* for the mRNA and promoter sequences associated with each transcript, comparing a range of *k*-mer lengths (see Methods; Glossary). We trained a series of ML classifiers to predict if a transcript was circadian or non-circadian in a binary classification system using the derived *k*-mer profiles for the same set of transcripts and MetaCycle derived labels used previously (for the transcriptomic ML model). Across the range of *k*-mers the best models were consistently generated with the classifier LightGBM and the most accurate model used a *k*-mer length of 6 to generate separate feature sets for the promoter and mRNA regions (8,192 features of *k*-mer counts per transcript) that were both inputted into the model (see Methods). This best optimized model showed (Figure 3a, Table S2): a mean F1 score of 0.766 on cross validation (standard deviation 0.006) and a test F1 score of 0.751 on class 0 (non-circadian) and 0.804 on class 1 (circadian). Again, our accuracy was largely balanced between the classes. An optimal *k*-mer length of 6bp for this analysis could reflect this being the smallest length *k*-mer that we would not expect to simply occur by chance, therefore giving ideal resolution. Due to the large number of features created when using a *k*-mer length of 6, using feature selection we tested the accuracy of our rhythmic classification when subsets of the feature set were used (Figure 3b; Glossary). We can reduce the feature number to ~200 and still achieve an F1 score above 0.7, but the highest accuracy was achieved with all 8,192 features and as such, for downstream investigations we used the full feature set.

**Figure 3.**
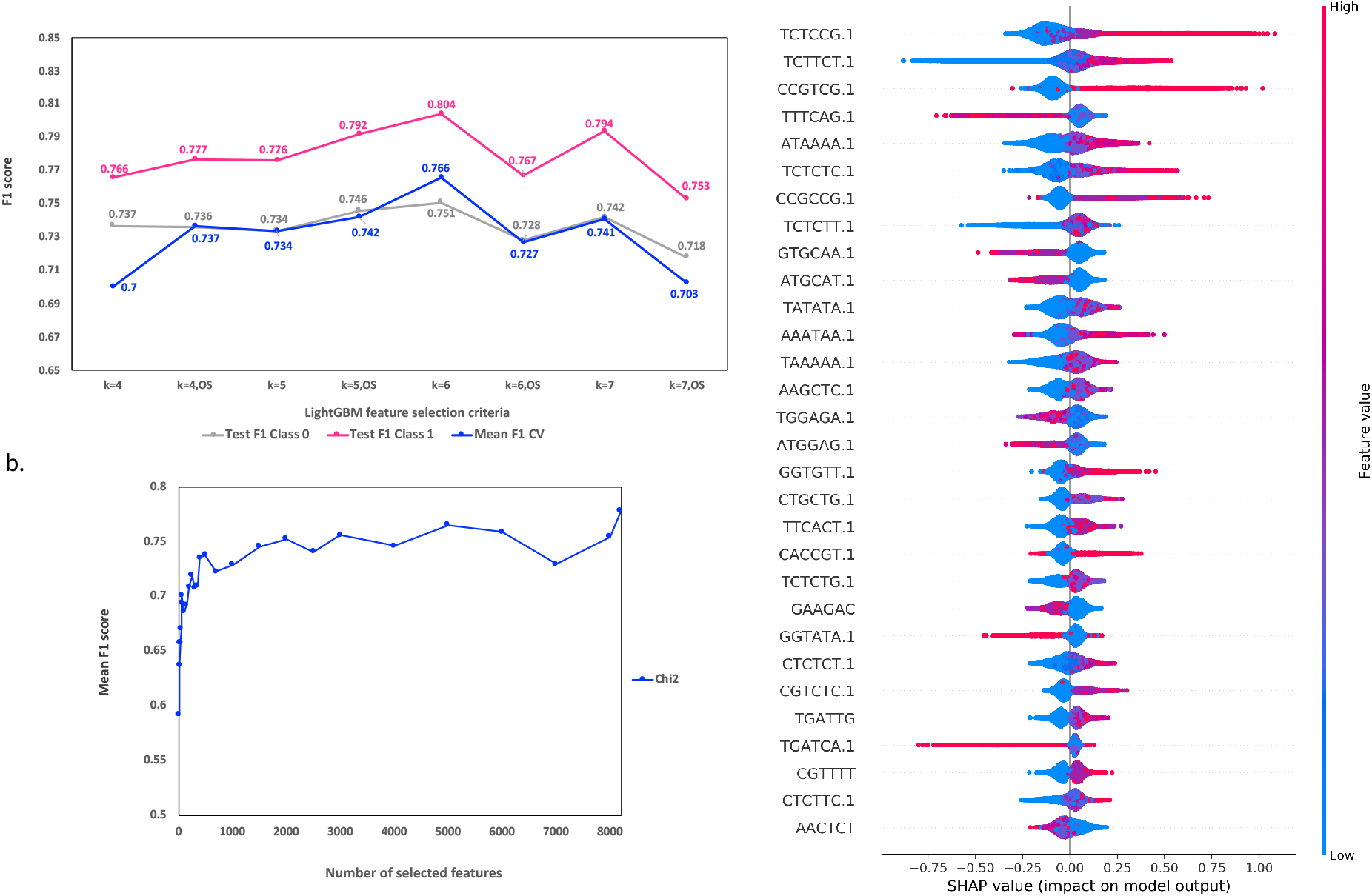
Arabidopsis circadian/non-circadian ML binary classification analysis using *k*-mer profiles. For our best performing classifier LightGBM we compare the F1 scores generated using **(a)** different *k*-mer lengths (4-7bp) for classification, with or without the use of oversampling (OS) since our classes are not perfectly balanced (Glossary). **(b)** To obtain each reduced set of *k-*mers we use chi-square (Chi2) feature selection. Here we show the best F1 score after 5-fold cross validation for each set of reduced features. **(c)** shows the top 30 most impactful features for predicting class 1 (circadian) considering all samples in the dataset (training and test) as calculated using SHAP (Shapley Additive exPlanations) (Glossary). Feature value equates to the frequency of a *k-*mer per transcript. When the frequency of a *k-*mer per transcript is high (red) and it has a positive SHAP value, this high frequency is driving the prediction of a circadian transcript. This is often coupled to the situation where the lower frequency of the same *k-*mer per transcript (blue) has a negative SHAP value, so the absence of the *k-*mer is driving the prediction of a non-circadian transcript. On the contrary, when the frequency of a *k-*mer per transcript is high (red) and has a negative SHAP value, the high frequency is driving the prediction of a non-circadian transcript. This is often coupled to the situation where the lower frequency of the *k-*mer per transcript (blue) has a positive SHAP value, so the absence of the *k-* mer is driving the prediction of a circadian transcript. Features e.g. the *k*-mer TATTGC, are labelled as “TATTGC” for counts from the promoter and “TATTGC .1” for counts from the mRNA.

Our *de-novo k*-mer generation approach allows downstream identification and investigation of both known and previously unknown sites with only the annotation of the TSS and TTS of a transcript required. Our short *k*-mers (6bp) should mainly represent regulatory elements such as TFBSs when derived from promoter/UTR regions. However, our inclusion of coding regions may allow us to encompass additional regulators e.g., miRNA binding sites. Although miRNAs tend to be 20-24bp in length, our *k*-mers may represent miRNA seed regions that are typically ~6bp in length and perfectly/near-perfectly match targets [17].

### Explanation of DNA sequence-based ML model links to circadian regulation

We next wanted to explain our model, to identify which *k-*mer’s were most influential in guiding it to predict transcripts as circadian, since these *k-*mer’s could represent the most critical regulatory elements for circadian regulation. If we observe known circadian regulatory elements in this process, this is also a means of validation of the model. As such, we used SHAP (Shapley Additive exPlanations) to explain our best DNA sequence-based model’s predictions by computing the contribution of each feature or *k-*mer to that prediction i.e., ranked feature impact on the classification (Glossary) [38]. We did this firstly at a global level by looking at the top 30 most impactful features across all of the transcripts for distinguishing class 1 (circadian) from class 0 (non-circadian) (Glossary; Figure 3c). Approximately half of the most impactful *k*-mers in Figure 3c show a positive correlation between *k*-mer frequency and the SHAP value or feature impact on the model. Higher frequencies of these *k*-mers for a transcript indicate a higher impact on it being classified as circadian. Of these positively correlated top 30 *k*-mers, 55% of those that contributed to the circadian classification of a transcript were predominantly in the promoter or the UTR of transcripts. We hypothesized that these *k-*mers represent TFBSs for transcription factors (TFs) linked to circadian regulation.

To investigate if our most impactful promoter/UTR *k-*mers for prediction were in fact TFBSs, we aligned known *Arabidopsis* TFBSs to each of the *k-*mers and filtered the most significant matches (Table S5; see Methods). We then validated the *k-*mers that match/likely represent TFBSs using experimental evidence or insight from the literature; many of the matched *k-*mers were closely associated with circadian regulation or circadian related processes. Notable *k*-mers of interest included (*k*-mer number 1; Table S5) matches to TFBS for two photo-responsive TFs (AT3G58630 and AT5G05550) (p-value 0.0002, e-value 0.18) which form interactions with a number of circadian-related proteins e.g. LIGHT INSENSITIVE PERIOD1 (LIP1), CONSTANS-Like (COL) 11 [39] and REVEILLE 2 (RVE2) [40]. Another *k*-mer (*k*-mer number 7; Table S5) matched a motif bound by several ethylene-responsive binding proteins (p=0.00003, e=0.02); ethylene synthesis is known to be both a circadian controlled process and also a moderator of the circadian clock [41,42]. We also found matches as would have been predicted for binding sites of known circadian TF’s including LUX ARRYTHMO (LUX) [43], CIRCADIAN CLOCK ASSOCIATED 1 (CCA1) [44] and LATE ELONGATED HYPOCOTYL (LHY) [45], alongside several motifs associated with light-induced or repressed sequences (SORLIP/SORLREP) (Table S5).

In contrast to our promoter/UTR *k*-mers, four of the positively correlated top 30 most impactful *k*-mer features defined by SHAP were observed primarily in coding regions across the circadian predicted transcripts. Since miRNAs are thought to influence circadian controlled processes [46,47] and are common in coding regions, we tested the possibility that these *k*-mers could represent miRNAs by aligning them (plus surrounding sequence) to mature ath-miRNA sequences to identify possible matches (see Methods). Two of the four *k-*mers and their flanking sequence matched miRNA sequences that were associated with developmental timing [48] and chloroplast biogenesis [49]. Therefore, for a subset of transcripts, the *k*-mers could represent putative miRNA binding sites that have been experimentally linked to circadian regulated processes, although this only accounts for a small proportion of the transcripts (Table S5). As such, we next investigated the possibility that these *k*-mers could represent RNA binding motifs (see Methods). In doing so we validated two of the *k*-mers by linking them to RNA binding motifs that are associated with circadian related processes. RNA-binding proteins are key regulators of gene expression and post-transcriptional regulation in eukaryotes, and, due to strong sequence conservation, their recognition preferences can be inferred from RNA-binding motifs. Two of the four coding sequence derived *k-*mers matched RNA-binding motifs (Table S5, p<0.05). The first is targeted by the RNA-binding protein Serine and Arginine Rich Splicing Factor 7 (SRSF7). This has been linked to circadian processes since circadian temperature cycles are known to drive rhythmic SR protein phosphorylation to control alternative splicing [50]. The *Arabidopsis* protein RSZ22 is a known true ortholog of the human SRSF7 SR factor that this alignment could represent [51]. The second *k-*mer matched motif is targeted by the RNA-binding protein LIN28A (*Homo sapiens).* The *Arabidopsis* protein Cold-Shock Protein 1 (CSP1) is a known homolog of LIN28A with a similar functional role in reprogramming, that this alignment could represent [52]. CSP1 has been implicated in seed germination timing that is also known to be clock related [53].

### Transcript-specific explanations reveal sub-classes within the binary class circadian

Our DNA sequence-based model used binary classification to discriminate transcripts under circadian regulation from those that are not, which is useful to identify circadian regulatory elements from model explanations. However, circadian rhythms reflect a variety of waveform shapes. As such, we bioinformatically identified co-expression modules (Glossary) from the transcriptomic profiles of the circadian transcripts that were used to train our ML models using weighted gene co-expression network analysis (WGCNA) [54]. This resulted in 8 modules with distinct circadian expression profiles. These modules represent groups of transcripts differentiated by phase of expression with the following observed (Figure S2); morning phases 0 (cluster 7) and 4 (cluster 5/6), day phase 8 (cluster 3), day/evening phase 12 (cluster 2), evening phase 16 (cluster 1) and night phase 20 (cluster 4/8).

We next sought to group our circadian transcripts into subgroups representative of different phases of expression, but rather than using transcriptomic information, this time we wanted to use the SHAP impact values of their *k*-mers. This effectively divides our DNA sequence-based model’s binary class circadian into multiple sub-classes, providing further insight into transcript rhythmicity. To enable this, we used model explanation of our best DNA sequence-based predictive model, but rather than identifying the most impactful *k*-mers in general (global explanation) for predicting class 1 (circadian), as previously, we now identify the most impactful *k*-mers for the classification of each circadian transcript individually (local explanation) (Glossary). For this, we focus on the true positive circadian transcripts where MetaCycle and our ML model predict circadian. These local explanations are transcript specific and could highlight *k*-mers that are regulating each transcript’s expression. Each transcript has a calculated SHAP impact value per feature (8,192 *k*-mers) and this set of values we refer to as the SHAP value profile for a transcript. The *k*-mer with the highest SHAP value being the most influential on the transcript’s classification as circadian. Comparison of these profiles allows us to compare and subdivide the transcripts within the binary class circadian, using DNA sequence composition related to gene regulation, rather than transcriptomic profile.

To investigate this, after deriving local explanations, we filtered the most circadian transcripts according to their SHAP explanation (“most positive cumulative SHAP value”, Figure 4a, see methods, Glossary). Then we focused on known circadian genes that were within this set i.e., experimentally validated and widely known true positive genes from previous studies. We clustered the derivative transcripts of these genes based on the similarity of their SHAP value profiles, which represent the relative impact of the *k*-mers on their classification as circadian (Figure 4b). In groups to the right of the dendrogram (purple), 85% of transcripts peak in their expression in the morning/day, whereas in groups to the left, 77% of transcripts peak in the evening/night (phases determined by MetaCycle). Therefore, circadian transcripts with more similar *k*-mer SHAP value profiles also had similar expression phases, thus dividing our circadian class into sub-classes representing phases of rhythmicity using *k*-mer information. For example, *PRR3* and *LUX* were found to have similar SHAP value profiles and we validated this by observing their similar transcriptomic expression profiles, with evening phases of expression of ZT15 and ZT13 respectively. Notable exceptions include the two LNK genes which have a transcript expression profile which peaks in the morning but have SHAP profiles similar to evening and night expressed genes, with *LNK1* most closely linked to *TOC1*. This suggests that *LNK1/LNK2* may be regulated by a separate mechanism to that regulating other dawn expressed genes. In the morning/day cluster we also see the gene *TIC* which peaks at dusk in the transcriptomic data; previously, rhythmicity of *TIC* was not detected in whole seedlings whereas here, we confidently classify this transcript as circadian from aerial tissue (MetaCycle q=0.004). Previous work concluded that *TIC* functions in the late evening [55] but plays a role regulating *LHY* that is in the same morning/day cluster as *TIC*, this may explain its appearance here [56]. Finally, we also see the night gene *PHYB* in the morning/day cluster, this may be due to the additional presence of the closely related *PHYA* in this cluster [57].

**Figure 4.**
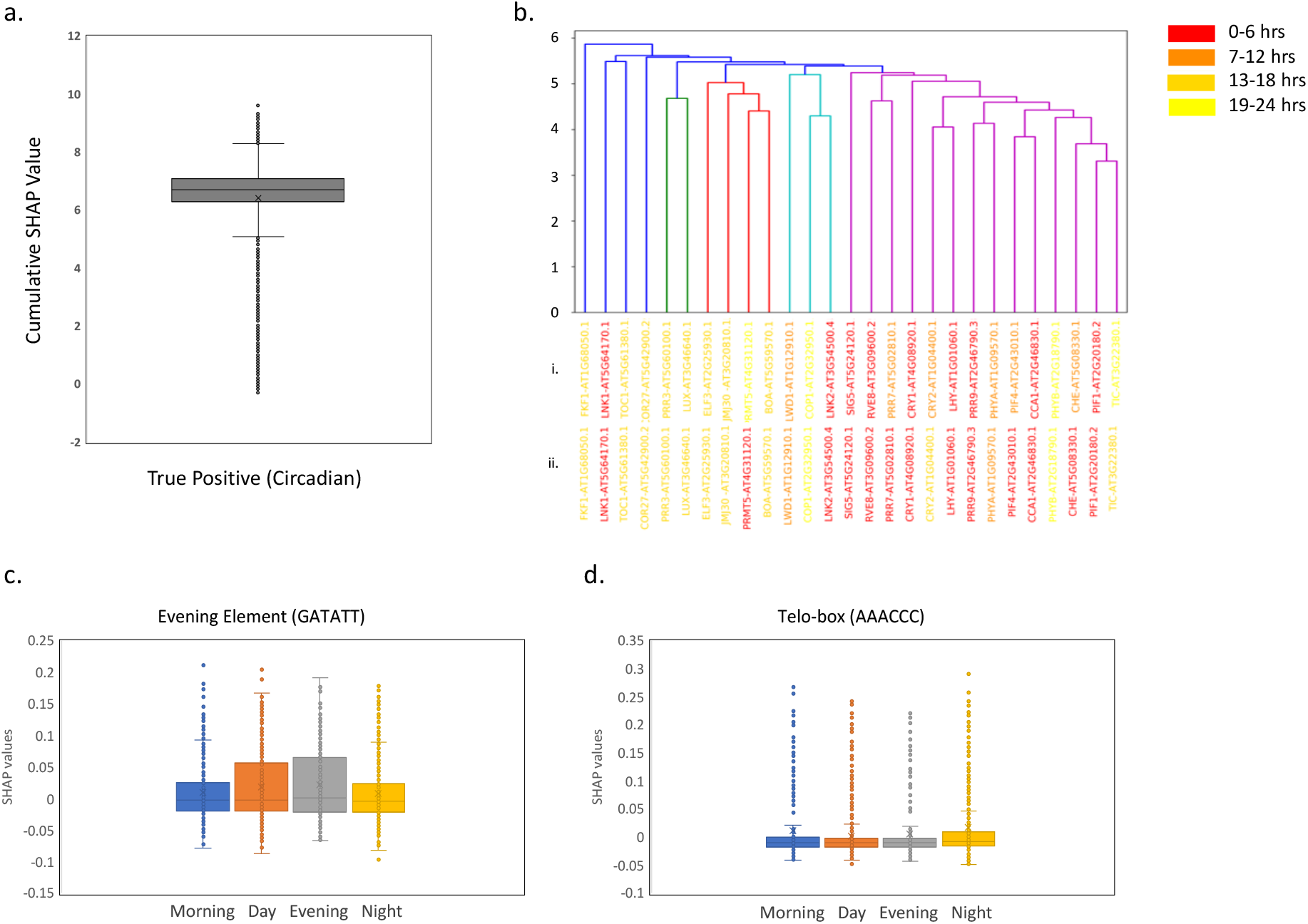
Investigating Arabidopsis circadian true positive transcripts after ML binary *k*-mer DNA sequence-based classification analysis. For our best performing classifier LightGBM. **(a)** Box plot to show the range of SHAP values across all true positive transcripts (correctly predicted as circadian). A positive SHAP value for a *k-*mer, for a specific transcript, indicates that the *k-*mer is driving the prediction of circadian, while a negative SHAP value indicates that the *k-*mer is driving the prediction of non-circadian for that transcript. SHAP values are summed for each transcript and the sum is defined here as the cumulative SHAP value. **(b)** Dendrogram produced by clustering known core circadian transcripts according to their profiles of SHAP values if the transcripts were also present in Q1-3 of **(a)**. We clustered transcripts using hierarchical clustering with average linkage and Euclidean distance (see Methods). Dendrogram labels coloured according to peak phases of expression; morning (0-6 hours), day (7-12 hours), evening (13-18 hours) and night (19-24 hours) as determined by **(i)** MetaCycle or **(ii)** the module of origin of the transcript from our 8 WGCNA generated modules. **(c-d)** Box plot to show the range of SHAP values across all true positive transcripts in groups morning day evening night for the specific *k*-mers **(c)** GATATT (Evening element) and **(d)** AAACCC (Telo-box).

We noted from our transcript SHAP value profile clustering (Figure 4b), that for sub-classes of transcripts with similar expression phases, the most impactful *k*-mers per sub-class could represent sequences that are regulating time-of-day specific expression. Identifying these using model explanation could facilitate the estimation of circadian expression phase without the need for a transcriptomic time course. To test this hypothesis, we split the transcripts into morning, day, evening and night and investigated which *k*-mers differentiated the groups. We identified the top 30 most variable *k*-mers between the four groups’ consensus SHAP explanations, these *k*-mers should therefore vary most in their impact between the groups (see Methods) (Table S6). Since we are comparing the *k*-mers that differentiate groups of transcripts that are separated by their phase of expression, we validated our hypothesis by matching the *k*-mers to binding sites that have been experimentally associated with specific times of day. For example, the late-night specific telo box [58], a G-box related sequence thought to associate with late night and dawn genes [59] and the Evening Element (EE) that appeared twice in the top 30 with two *k*-mers matching it. When we compared the importance of these *k*-mers between the morning, day, evening and night groups, the EE had a higher impact on model prediction in the evening group than in the other three groups and this difference was statistically significant compared to both morning and night (Figure 4c, Table S7). Additionally, the Telo-box had a higher impact on model prediction when observed in the night group compared to all other groups and this difference was statistically significant compared to day and evening, fitting with its late-night specificity (Figure 4d, Table S7).

### Case study: transcript-specific explanation for PHYA-E guides re-classification of PHYC

The PHYTOCHROME (*PHY*) genes encode red and far-red photoreceptors directly involved in setting the clock. Previous studies have identified circadian regulation of PHY A-E as rhythmic. [60]. However, PHYC/PHYD/PHYE were all called non-circadian by MetaCycle with q-values of 0.99, 0.60 and 0.13 respectively. These genes should be rhythmic, but this may not be clearly reflected in the transcriptomic data, likely due to their low amplitude expression patterns (Figure S3a). As a result, these genes were missing from downstream analysis and can be used as a case study of unseen test datapoints (Glossary) for the ML models. For the PHYA-E primary transcripts, Table S8 highlights MetaCycle’s 40% accuracy, only classifying PHYA-B as circadian, compared to our ML (12 timepoint) model’s 80% accuracy since we additionally classify PHYD-E as circadian. This is supported by visually evident rhythmic expression in the transcriptomic data, particularly for PHYE and to a lesser extent for PHYD (Figure 2a). We maintain our 80% accuracy when we generate *k-*mer profiles for the PHYA-E transcripts and use our DNA sequence or *k-*mer based ML model to predict circadian/non-circadian. Both of our ML models (transcriptomic and DNA-sequence-based) classify PHYC as non-circadian with the other primary PHY transcripts predicted circadian. Even the DNA sequence-based ML model discriminated PHYC from the other PHY transcripts despite sequence similarity between them. Moreover, the transcriptomic expression profile for PHYC provides an unconvincing circadian rhythm, with an amplitude tending towards zero (0.02), compared to the other transcripts (Figure S3a). Here, we assumed that all of the PHYA-E primary transcripts were circadian. This may reflect previous work that concluded a weak rhythmic association of PHYC potentially due to post-transcriptional circadian regulation not promoter regulated expression [60,61].

We used the SHAP explanations for the PHYA-E transcripts to identify the regulatory elements that were most impactful in guiding their classifications, using the DNA sequence-based model. We compared the SHAP impact values between each of the PHY transcripts A/B/D/E (circadian) and PHYC (non-circadian) to identify those *k-*mers or regulatory elements that are most impactful in predicting PHYA/B/D/E to be circadian but also in predicting PHYC to be non-circadian (six identified in Table S9). The change in frequency of these *k-*mers is most likely to be responsible for the circadian/non-circadian predictive differences between the transcripts according to our model (Supplementary Note 3; Figure 5). To investigate if altering any of the six identified *k*-mers (Table S9) had the potential to induce rhythmicity in PHYC, we sequentially evolved the spectrum of PHYC, one *k*-mer at a time, to mimic the robustly rhythmic PHYA/B transcripts more and more with each iteration. We used our DNA sequence-based ML model to classify the evolved transcripts. Firstly, removing *k*-mers GGTAGA then TTTCTG sites, resulted in predictive probabilities for the circadian class of 0.42 and 0.48 respectively (increasing from 0.38). Secondly, adding AAATAA increased the predictive probability of circadian class membership further to 0.58. Finally, adding TCTCCG resulted in a circadian class predictive probability of 0.75 and placed this transcript’s classification now confidently as circadian. We noted that some potential regulatory elements are more important than others, having a larger effect on the classification of the transcript; for example, *k*-mers in the 5’UTR had a larger effect on classification. Additionally, we show that multiple elements combine to have a greater impact on transcript classification and potentially regulation.

**Figure 5.**
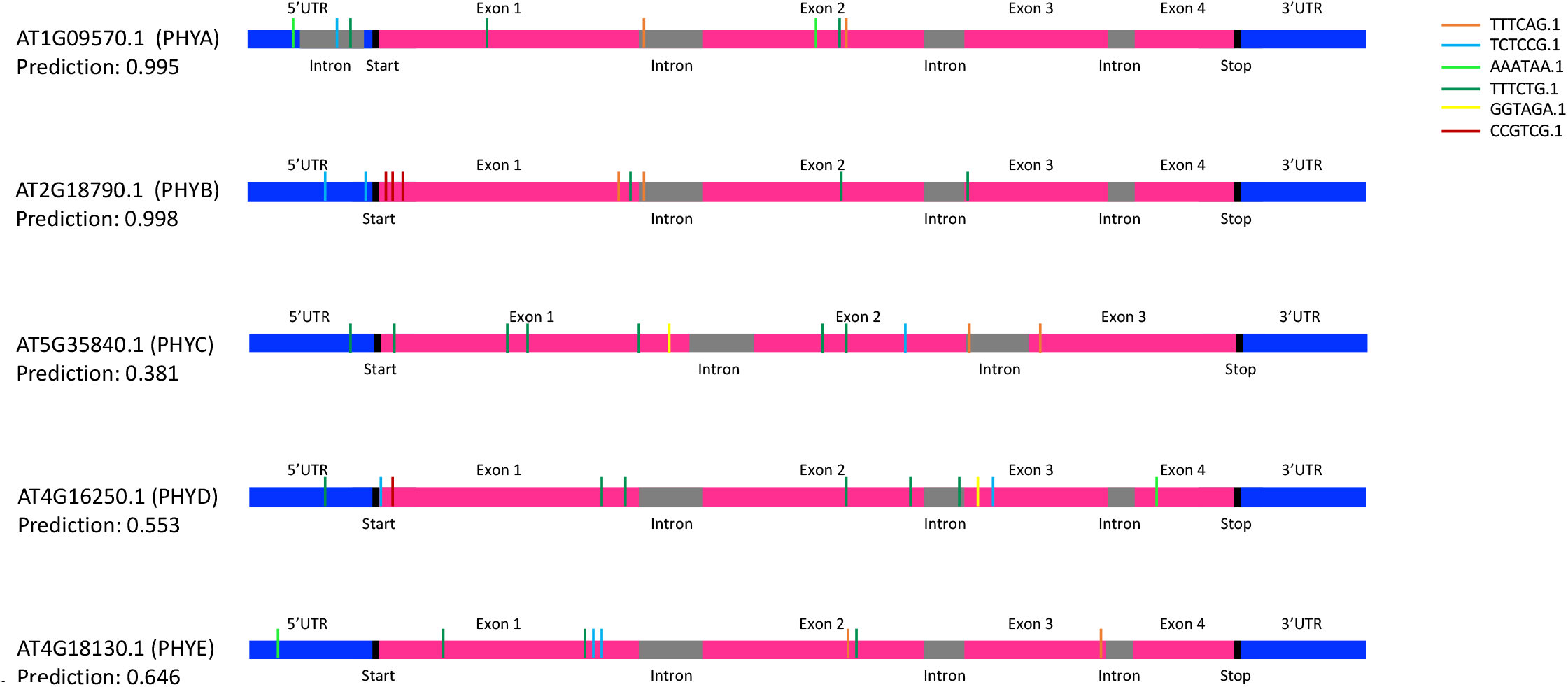
Investigating Arabidopsis PHYA-E transcripts after ML binary *k*-mer classification analysis. We compared the SHAP explanations between each of the primary PHY transcripts A/B/D/E and PHYC. Here a high comparative number translates to regulatory elements being more impactful in predicting PHY A/B/D/E to be circadian but also typically more impactful in predicting PHYC to be non-circadian. Schematics of the transcript sequences for PHYA-E and the associated start positions of the 6bp k-mers TTTCAG, TCTCCG, AAATAA, TTTCTG, GGTAGA and CCGTCG, that were identified as in the top three highest values per comparison (PHYA versus PHYC, PHYB versus PHYC, PHYD versus PHYC and PHYE versus PHYC) with the largest differences in SHAP values (Table S9; Supplementary Note 3).

We aligned known *Arabidopsis* TFBSs to the UTR-based *k-*mers from PHYA/B that most positively impacted PHYCs circadian re-classification during our evolution to suggest biological reasons why these sites may be having such a large effect. Firstly, AAATAA aligned to the TFBS of MYB56 that is involved in the regulation of anthocyanin levels in response to circadian rhythms [62] (Table S5). Secondly, TCTCCG matched TFBS of AT3G58630 that has a protein-protein interaction with LIP1 a gene known to function in the clock regulating light input downstream of photoreceptors such as PHYB [63].

To extend this analysis beyond the well-known PHYA-E genes, we collated a further 41 known key circadian genes, with published evidence of rhythmic expression from across the literature and compared the classification accuracy of their associated primary transcripts between MetaCycle, our ML model using 12 timepoints and our ML model using DNA sequence (Table S10). MetaCycle shows an overall accuracy of 80.49% classifying the 41 transcripts as circadian compared to 95.12% with the ML transcriptomic model (Table S11). We tested 10 of the 41 genes that were not used to train either of our ML models and were therefore unseen datapoints, mainly due to MetaCycle not assigning a highly confident classification to their transcripts (q<0.01) due to low amplitude expression profiles. These are problematic transcripts for classification and can be used as a measure of the worst-case scenario for predictions. Using 12 timepoints our ML model was much more accurate at correctly classifying these transcripts as circadian despite their problematic low amplitude rhythms (80% accuracy versus 20% for MetaCycle). This suggests that our model has the potential to generalize well to unseen transcripts. Interestingly, our model that used DNA sequence alone achieved a higher accuracy of 90% on the unseen datapoints, which was much closer to its recorded accuracy on all 41 genes (92.68%) sidestepping the problems associated with low recorded amplitudes using genetic sequence features.

### Predictions using DNA sequence generalize to other *Arabidopsis* ecotypes

We previously ascertained that our ML model (using DNA sequence) can accurately make predictions on unseen datapoints. We assessed this in both our initial testing (with held out test data; Glossary) and in our case study analysis of known circadian genes. We next want to assess how well our model performs on unseen DNA from a different source to that used for model training (Col-0). We selected the *Arabidopsis* ecotype Ws-2 for this test, generating *k-*mer spectra for related transcripts and using the transcriptomic dataset generated by [10] to label Ws-2 transcripts circadian/non-circadian to gauge accuracy (Table S1; Supplementary Note 4; Figure S4). From this analysis, 71.4% of Ws-2 DNA sequence-based classifications matched their labels derived from [10] transcriptomic data. This is only ~5% lower than the accuracy given by the DNA sequence-based model using Col-0 (mean F1 score of 0.766 on cross validation) and therefore, we see only a minimal decrease in accuracy applying our model to a new ecotype (Supplementary Note 5).

We next wanted to use our DNA sequence-based model to identify transcripts that differentiated in rhythmicity between *Arabidopsis* ecotypes. Then we use model explanation to explain which regulatory elements influence this and can validate findings. Such functionality gives tremendous power for downstream gene expression manipulation. We identified 12 transcripts that were classified as circadian for Col-0 but non-circadian for Ws-2 by the DNA sequence-based model (both with a predictive probability >0.8) (Table S12). We ranked the transcripts according to the predictive probability of them being circadian for Col-0 and the corresponding predictive probability of them being non-circadian for Ws-2. Our most confident or top ranked transcript was AT1G58602.1-RECOGNITION OF PERONOSPORA PARASITICA 7 (*RPP7*) i.e. the most probable circadian transcript in Col-0 (probability 0.999) and the most probable non-circadian in Ws-2 (probability 0.991). RPP genes have been previously reported to confer resistance to races of *P. parasitica* in an ecotype specific manner. A functional copy of RPP7 is thought to mediate resistance to infection by the *Peronospora* isolate Hiks1. Work by [64] found that while Col-0 has a functional RPP7 and is resistant to Hiks1, Ws-2 is susceptible to attack by this pathogen. This coincides with our DNA-sequence predictions suggesting that the circadian behaviour of RPP7 is important for defence functionality. This conclusion is also supported in the experimental transcriptomic data where *RPP7* in Ws-2 shows consistent low expression but in Col-0 it is expressed at higher levels with a circadian rhythm (Figure S5a) [64]. RPP7 has been linked to circadian regulation; firstly, because resistance (R)-genes in the RPP family were reported to be under CCA1 control [65], and secondly, via RPP7’s required interactor EDM2 that is involved in the promotion of floral transition by regulating the floral repressor *FLC* [66].

Previous evidence supports our observed differentiation in rhythmicity of RPP7 between Col-0 and Ws-2. However, our advantage would be to use model explanation to understand which elements differ between Col-0 and Ws-2; in this example in Ws-2, this could represent which elements to change to render it resistant to Hiks2. As such, for each k-mer, we compared the SHAP impact values from the DNA sequence-based model between the Col-0 and Ws-2 homologs of AT1G58602.1 (*RPP7)*. We ranked the *k*-mers in ascending order as the difference in SHAP impact values between the homologs increased, to highlight the regulatory elements that were most impactful in guiding the differential circadian/non-circadian predictions (Figure S5). The top 5 ranked *k*-mers, according to differences in SHAP impact, closely linked either to the circadian clock or to disease resistance mechanisms, or both (Supplementary Note 6). We then sequentially evolved the *k*-mer spectrum for AT1G58602.1 in Ws-2, a *k*-mer at a time to match Col-0 more and more with each iteration. Each iterative evolved transcript was classified using the DNA sequence-based model, where we observed that the predictive probability of the circadian class for each evolved gene quickly increased (Figure S5b). Adaptation of 26 Ws-2 *k*-mers to match Col-0 was needed to change the prediction for Ws-2 from non-circadian to circadian and adaptation of 124 Ws-2 *k*-mers was needed to reach the maximum predictive probability of 0.999. We noted that the predictive probability of the circadian class for Ws-2 was highly positively correlated (0.676) with the difference in SHAP values between the Col-0 and Ws-2 *k*-mers (Figure S5c). Our analysis shows that some regulatory elements have a larger effect on the classification of the transcript than others and that this effect is quantifiable using model explanation. We also show the potential for large combinations of regulatory elements to work together, potentially each contributing a small amount, to result in a large overall impact on gene classification and potentially regulation e.g., the 26 *k*-mers that we changed here to convert Ws-2 to be classified as circadian.

### Identifying a set of transcriptional biomarkers that predict internal circadian time

To complete our suite of circadian resources, here, as opposed to identifying transcript rhythmicity, we consider the experiment as a whole, using ML to determine the circadian time of sampling i.e. predicting the phase of the endogenous circadian clock, using a set of transcriptional biomarkers from any single transcriptomic timepoint. Previous studies have developed such models for human and mammalian transcriptome data sets [32,33,34,67,68]. However, we develop a new method that we apply to plant data that innovatively uses model interpretation to identify a set of new *Arabidopsis* biomarker transcripts to guide our predictions. This incorporates, biomarker selection from across circadian phases to increase accuracy and robustness.

To train our model we used the TPM normalised circadian dataset described earlier [8] and the two further transcriptomic datasets [9,10] for validation and testing (see methods; Glossary). Firstly, we aggregated a selection of metrics to rank and select transcript subsets from [8] according to their confidence of rhythmicity for model training. Table S13 highlights the mean absolute errors (MAE) of the predictions of circadian time without hyperparameter optimization (Glossary) on the three temporal transcriptomic datasets, using different sized subsets of the highest ranked rhythmic genes. The lowest MAE, based on the [10] test dataset, was 104 minutes and was observed with a selected subset of 50 transcripts. Using confidence of rhythmicity for transcript prioritization, we noted that the representation of our subsets of transcripts across the 8 co-expression modules generated by the WGCNA gene co-expression network analysis was not uniform (Figure S6a; Glossary). This reflects an uneven representation across the phases of rhythmic expression. Therefore, secondly, we prioritized selection of transcripts using model interpretation in the form of feature selection to make the frequency distribution across the modules more uniform (see Methods; Glossary). Optimizing performance based on the validation dataset, our best performing model overall used a final subset of 15 transcripts (Table S14) and had a MAE of 21 minutes on the training data, 56 minutes on the [9] validation data and 46 minutes on the test data from [10]. Figure S6b and S6c also highlight that after such feature selection there was a decrease in the generalisation error on average across the [10] test dataset with the improvements in MAE decreasing as the number of genes increased. This supports the theory that features containing different temporal patterns of varying strengths outperform features containing strong but highly correlated patterns.

The performance of our best model (15 transcripts with a MAE of 46 minutes on the test data) is in line with the ~1-hour test error reported by [67] using their state-of-the-art method ZeitZeiger. As such, we applied ZeitZeiger to our datasets [8,9,10], to compare directly with our model. To reflect our previous approach, firstly, dataset [8] was used to fit ZeitZeiger, with predictions then being generated on the validation [9] and testing [10] datasets to compare with the predictions generated by our method. Our approach significantly outperformed ZeitZeiger on the test dataset (MAE of 46 compared to 143 minutes, Figure S7) demonstrating our efficacy at generating highly accurate predictions for circadian time. We also noted a large disparity in training, validation and test errors by ZeitZeiger (MAE of 6 minutes on training, 119 on validation and 143 on test) that suggests overfitting (Glossary). We hypothesized that our selection of biomarker transcripts to ensure even representation across the phases of rhythmic expression, would yield a more robust or generalizable mapping from expression data to internal circadian time i.e., less overfitting; this analysis supports this hypothesis.

The 15 transcripts in our final subset act as a small subgroup of biomarker transcripts that are sufficient to allow prediction of the circadian time (Table S14). Interestingly, the 15 transcripts did not include any core clock genes. This analysis was conducted using the ecotype Col-0. However, using the Ws-2 data [10] a MAE on this ecotype of only 53 minutes was observed (5 minutes lower than for Col-0 on which the model was trained). Generally, we observed no relationship between circadian time and prediction error except for in the training dataset where errors at the 20-hour timepoint were significantly larger than the other times (Figure S6d). However, variation in error across the timepoints typically stayed under 90 minutes allowing sufficient resolution of circadian time given that typical sampling strategies are between 2-4 hourly.

## Conclusions

We describe a series of ML based approaches that enable cost-effective analysis and insight into circadian regulation in *Arabidopsis*. One of the drawbacks of ML is a lack of clarity as to why it makes specific predictions. We focus on illuminating what is inside the ‘black box’ via explanation or interpretation of predictive ML models. Although we demonstrate this for circadian rhythms, this approach has widespread implications for other complex or temporal gene expression patterns.

When we predict circadian transcripts using low numbers of mRNA-seq timepoints, not only do we improve accuracy compared to existing methods, but we also use model interpretation to optimize sampling strategies. Some of the most accurate reduced sampling strategies that we identify align with existing approaches e.g., timepoints spaced evenly across a day. However, other identified strategies were unexpected e.g., consecutive timepoints or those across the intersect of day 1 and 2.

Most significantly, we use *only* DNA sequence features for accurate circadian classification, requiring no prior knowledge of regulatory elements or transcriptomic data. This offers advantages over existing methods to not only predict expression but to decipher regulation at the same time since, using an explainable AI algorithm, we define regulatory elements on the fly as we make predictions. Automated definition and prioritization of these feature profiles for transcripts, *de novo*, using AI, has the potential to support functional annotation of genomes and precision agriculture. This application could re-define how we generate testable hypotheses to understand gene expression control. Our predictive accuracy is possibly higher than our current estimates as our DNA based approach scores the potential of a gene to be circadian regulated. However, it is possible that this regulation may be restricted to specific tissue types or developmental stages. Therefore, our experimental generated labels may be underestimating the number of rhythmic genes. We propose incorporating both DNA sequence features with epigenetic or additional biological features into predictive models to refine predictions, since epigenetic modifications are thought to effect tissue-specific gene expression.

Finally, we predict circadian time while using model interpretation to derive novel *Arabidopsis* marker transcripts. These selected transcripts could be used to test single datapoints in existing and emerging *Arabidopsis* datasets to investigate how genotypes, treatments and environmental conditions affect circadian clock function.

## Glossary

Machine Learning (ML): A branch of artificial intelligence based on the idea that systems can learn from data, identify patterns and make decisions with minimal human intervention.
Model interpretation/explanation/explainable AI: A set of methods and algorithms that help us to understand and interpret the predictions made by ML models.
Features e.g. DNA sequence features: Input variables into ML models, a feature is a measurable property or characteristic related to the phenotype being observed/predicted.
ML classifiers: A ML classifier is an algorithm that predicts the class (e.g., circadian or non-circadian) of given data points (e.g., transcripts). A ML classifier utilizes training data to understand how given input variables or features of a data point relate to a specific class or classes. Once the classifier is trained, it can predict the class for unseen data points in the test data.
Binary classification system: Classification is a supervised learning approach in which the ML model learns from the input data or feature set and then uses this learning to classify new observations into one of two possible classes (e.g., circadian or non-circadian).
Training data: The data used to train a ML algorithm or model e.g., a table where the rows are the data points such as transcripts and the columns are the features describing the data points.
Test data (held out): A dataset (e.g., a table transcripts x features) that is independent of the training dataset, the model has not seen this data during training-it is “held out”. If a ML model has been fitted to the training dataset and then also fits the test dataset well (shows accurate predictive performance on the test dataset) then we would say that minimal overfitting has taken place.
Model validation: An independent dataset (e.g., a table transcripts x features) that is specifically used to tune the parameters of a ML classifier i.e. used to measure performance and guide model training.
LightGBM: Short for Light Gradient Boosting Machine, is a distributed gradient boosting framework for ML. It is based on decision tree algorithms and used for ranking, classification and other ML tasks e.g., the LightGBM classifier is used for classification tasks.
Cross Validation (CV): Cross Validation is a technique that is used to evaluate ML models. It involves training several ML models on subsets of the available input data (also known as folds) and evaluating them on the remaining (held out) subset of the data. For example, in k-fold cross-validation, you split the input data into k subsets or folds of data specifically.
Parameters or Hyper-parameters: The part of the ML model (e.g., LightGBM) that is learned from the training data. If the ML model is the hypothesis then the parameters are used to tailor the hypothesis to the training data.
Fine tuning: Making small adjustments to the hyper-parameters of a ML model using the training data while performing CV to achieve the desired output or optimized performance (here higher accuracy).
True positive (TP): The model correctly predicts the positive class or class 1 (e.g., circadian)
True negative (TN): The model correctly predicts the negative class or class 0 (e.g., non-circadian).
False positive (FP): The model incorrectly predicts the positive class (e.g., predicts circadian but the true class is non-circadian).
False negative (FN): The model incorrectly predicts the negative class (e.g., predicts non-circadian but the true class is circadian).
Confusion Matrix: A table/matrix with two rows and two columns that reports the number of false positives, false negatives, true positives, and true negatives computed from a ML model’s prediction on a subset of data e.g. test, training or both.
Precision (P): The ratio of correctly predicted positive observations (true positives) to the total predicted positive observation. P=TP/(TP+FP). Precision answers the question “Of all the predicted positive observations (e.g., correctly predicted circadian), how many were actually positives (e.g., true circadian)?”. High precision score relates to low false positive rates.
Recall (R): The ratio of the correctly predicted positive observations to the total of observations in the positive class. R=TP/(TP+FN). Recall answers the question “Of all the true positive observations (e.g., true circadian), how many were correctly predicted as positive (e.g., circadian) by the model?”
F1 score: The F1-score is a measure of the accuracy of a ML model. More precisely, the F1-score is the weighted average (harmonic mean) of calculated from the precision and recall. Note that usually when precision increases, recall decreases and vice versa. The highest possible F1 score is 1, indicating perfect precision and recall, and the lowest possible value is 0.
Feature selection: These methods are used to reduce the number of input variables or features to those that are thought to be most useful to a ML model in order to predict the target variable (here circadian/non-circadian class).
Eli5 Feature selection: Computes feature importance for any ML model by measuring how, in this case, the F1 accuracy score decreases when a feature is not available. This method is also known as “permutation importance” or “Mean Decrease Accuracy (MDA)”.
Chi2 Feature selection: Can be used to measure the dependence between input features to the ML model and class. This score can be used to select the features with the highest values for the test chi-squared statistic, relative to the classes. Using this function “weeds out” the features that are the most likely to be independent of class and therefore irrelevant for classification.
Generalizable: The ability of a ML model to maintain its accuracy across a range of different datasets e.g., here we apply a model trained on *Arabidopsis* to the divergent species wheat.
*k*-mer profiles: Sub-sequences of length *k*, composed of nucleotides (A, T, G, and C) and contained within a biological DNA sequence. Our selected *k*-mer length of 6 yields 4^6^ (4096) possible *k-*mers from the 4 nucleotides in the DNA alphabet. Every possible *k*-mer is counted in each transcript and promoter (separate counts) generating a *k*-mer count numerical profile of 4096*2 = 8192 features.
Oversampling: Involves randomly selecting samples from the minority class (class with lower numbers of training samples), typically replicating them and adding them to the training dataset to even the number of samples between the classes. Rather than replicating the minority observations/samples it is alternatively possible to create synthetic observations based upon the existing minority observations.
SHAP: An explainable AI algorithm, called Shapley Additive exPlanations - SHAP. SHAP combines game theory with local explanation enabling accurate interpretations on why and how a ML model predicted a particular value (in our case a binary class) for a given sample.
SHAP impact values: For binary classification using our DNA sequence-based model, the SHAP explainer returns two SHAP value tables (transcripts x k-mer-based features), one for the class 0 (non-circadian) and one for the class 1 (circadian). These SHAP values represent the contribution of each feature to that prediction i.e. ranked feature impact on the transcript classification distinguishing class 1 (circadian) from class 0 (non-circadian).
Global explanation: Looking at the most impactful features across all of the transcripts for distinguishing class 1 (circadian) from class 0 (non-circadian).
Co-expression modules: Correspond to clusters of genes that have a similar shape expression profile across the transcriptomic time series. They are likely to have similar functions or involve common biological processes.
Local explanation: Rather than identifying the most impactful features in general (global explanation) for predicting class 1 (circadian) across all of the transcripts, distinctly, local explanation relates to the identification of the most impactful features for the classification of each circadian transcript individually i.e., transcript-specific explanations. A positive SHAP value for a feature, for a specific transcript, indicates that the feature is driving the prediction of circadian, while a negative SHAP value indicates that the feature is driving the prediction of non-circadian for that transcript.
Cumulative SHAP value: SHAP values are summed for each transcript and the sum is defined here as the cumulative SHAP value.

## Methods

### Data generation

The datasets used in this analysis are detailed in Table S1. All previously published datasets have details for data generation in the relevant associated publication. For the wheat time course: Cadenza seedlings were grown under 12:12 light:dark cycles at 22C for 14 days before transfer to constant light. After 24 hours under constant conditions, whole aerial tissue samples were taken every 2 hours for 3 days starting at perceived dawn (ZT=0). Total RNA was extracted using Qiagen RNeasy plant mini kits. Illumina TruSeq strand specific libraries and mRNA-seq was carried out by Novogene Co. Ltd. 150bp PE reads were generated from each library to an average depth of 70M reads.

### Bioinformatic analysis of transcriptomic information

Arabidopsis: Raw reads were obtained in FASTQ format for each Arabidopsis dataset [8,9,10]. These reads were filtered for quality, and any remaining adaptor sequence trimmed with Trimmomatic [69]. Surviving reads were aligned to the *Arabidopsis thaliana* genome (TAIR 10) using HISAT2 [70] with default parameters, except for maximum intron length, which was set at 5000nt. Uniquely mapped transcripts were quantified using StringTie [71] and the raw expression counts per transcript, for each replicate were subsequently normalised using DESeq2 [72]. A custom Perl script was also used to extract the TPM values from StringTie quantifications.

Wheat: The wheat mRNA-seq samples, of 150bp PE reads were aligned, quantified and normalised as described above, except that HISAT2 was used with default parameters and reads were mapped to the Chinese Spring RefSeq v1.0 wheat genome [73].

### Defining circadian genes using MetaCycle

Initially, Metacycle [13] was implemented on the [8] *Arabidopsis DESeq2* normalized gene expression counts (average expression count across two biological replicates, per transcript) to classify rhythmic expression using the 12 timepoints. This analysis included 44,963 transcripts. Metacycle (meta2d) was run with the following parameters on each of the normalized and non-normalized datasets; minimum period length of 18, maximum period length of 30, ARSER/JTK-CYCLE/Lomb-Scargle methods used [35–37], phase adjustment with predicted period length and the Fishers method was used to integrate multiple P-values. The output of this analysis includes a measure of period (the integrated period from MetaCycle is an arithmetic mean value of multiple periods from the implemented methods), phase (phase integration is based on the mean of circular quantities) and amplitude (amplitude is associated with general expression level and relative amplitude is used to compare amplitudes of genes with different expression levels). Finally, Benjamini-Hochberg q-values (BH.Q) were reported. Typically, significantly rhythmic gene expression profiles are defined at values q<0.05, we also use q<0.02 to limit selections based on the highest confidence.

MetaCycle was also implemented on wheat (variety Cadenza) transcriptomic timepoints (Table S1). Here, we also used DESeq2 normalized gene expression counts (average expression count across four biological replicates, per transcript) to classify rhythmic expression using the 24 timepoints. MetaCycle was used to detect rhythmicity in the normalized time course dataset with the same parameters used previously for *Arabidopsis*. MetaCycle classified 30,065 out of 112,955 analysed high confidence transcripts as circadian using a maximum q-value of 0.05. To select wheat transcripts as a test dataset for the *Arabidopsis* Col-0 trained transcriptomic ML model, we focused on 25,000 transcripts that MetaCycle classified (labelled) as highly circadian i.e. with high confidence (q<0.015) and 25,000 of the least likely candidates to be circadian genes (q>0.99) identified by MetaCycle with 24 timepoints.

### Clustering circadian transcripts according to transcriptomic profiles

Gene co-expression analysis was carried out using the R package WGCNA [54]. The 9,394 transcripts identified by MetaCycle as significantly rhythmic (q-value < 0.05) were filtered to remove transcripts where the sum of normalised expression counts across 21 or more replicates was less than 10. The remaining 8,136 transcripts were used to construct signed hybrid networks on a replicate basis using the blockwiseModules() function. The soft power threshold was calculated as 16, and the following parameters were used; minModuleSize = 30, corType = bicor, maxPOutliers = 0.05, mergeCutHeight = 0.15. Highly connected hub genes were identified for each of the eight co-expression modules using the function chooseTopHubInEachModule().

### Binary classification: ML model training and tuning

We used Scikit Learn (v3.7) for the ML binary classification analysis to predict if a gene was circadian or not with either transcriptomic or DNA sequence-based feature sets [74]. Unless otherwise stated, the MinMaxScaler was used to scale the features from 0 to 1, 90% of the data was used for training and the remaining 10% was held out for testing. 5-fold cross validation was performed on the training data. We used K-folds for cross validation (n_splits=5). The methods’ hyperparameters were optimized using a grid search to test a range of parameters (Table S15) for the following classifiers: Logistic Regression, Gaussian process, Random Forest, XGBoost, LightGBM, Support Vector Machine (SVM) (linear kernel), Decision Tree and K nearest neighbours (KNN). We selected the best ML model for each use-case (using best parameters after fine tuning), according to the highest F1-score on test set and cross validation.

The features used to train our initial transcriptomic (12 timepoint) ML model were the normalized averaged expression profiles for each *Arabidopsis* Col-0 transcript [8] and the “baseline gold-standard” circadian/non-circadian labels as defined by MetaCycle using 12 timepoints and stated in the main text and methods (Supplementary Note 1). We trained ML classifiers to predict if a transcript was circadian or non-circadian in a binary classification system using 7,734 of the least likely candidates to be circadian (q>0.99) labelled by MetaCycle alongside the 7,734 highly circadian transcripts (q<0.02). Additional transcriptomic models developed downstream, were trained using reduced numbers of timepoints either from the same [8] dataset or from a different data source (*Arabidopsis* Col-0 from [10]) with the same “baseline gold-standard” labels as previously. All transcriptomic models use normalized averaged expression profiles for each transcript.

To test the accuracy of our best trained transcriptomic ML binary classification model that uses 3 timepoints; for *Arabidopsis* Col-0 [9] test data, we assessed all predictions with a prediction probability or confidence of 95% or more and expressed those classed correctly as a proportion of the correct plus incorrect predictions to gain an overall accuracy percentage (for *Arabidopsis* this encompassed 14,652 predictions). Since the [9] test data is derived from *Arabidopsis* Col-0 we used the original [8] MetaCycle derived “baseline gold-standard” Col-0 labels to calculate accuracy. For wheat, we tested accuracy using the 50,000 genes (25,000 circadian and 25,000 non-circadian labelled by MetaCycle) that have already been filtered to encompass highly circadian and non-circadian representative genes, therefore, here we use the overall F1 score for our predictions directly. The features/attributes used to train our DNA sequence-based ML model were the *k-*mer profiles for each transcript (*Arabidopsis* Col-0) and circadian/non-circadian “baseline gold-standard” labels as defined by MetaCycle. To train our initial model, we generated *k*-mer profiles *de-novo* for the mRNA and promoter sequences associated with each transcript. We trained a series of ML classifiers to predict if a transcript was circadian or non-circadian in a binary classification system using 6,907 of the least likely candidates to be circadian alongside the 7,481 of the highly circadian transcripts used previously. However, these numbers were reduced from the 7,734 used previously due to our focus on mRNA only (removing ncRNA, snoRNA and lncRNA’s). To develop our feature sets we trialled *k*-mers from 4-7bp in length to encompass a range from smaller *k*-mers that we expect to see by chance to larger *k*-mers that we would not expect to see by chance in a promoter (1,500bp) or mRNA region (average length 2069bp); *k*-mers of 4, 5, 6, 7bp occur by chance every 256, 1024, 4096 and 16,384bp respectively, given equal frequency of the four nucleotides. Since we trialled *k*-mers ranging from 4-7bp in length, our feature numbers varied from 256-16,384 to reflect the numbers of possible *k*-mers (4^k^) from the 4 nucleotides in the DNA alphabet. Every possible *k*-mer of length k, e.g. 6bp, is counted in each transcript and promoter (separate counts) defined as 1500bp upstream of the TSS, meaning that no prior knowledge of regulatory elements or detailed gene annotation is required. Although we tested combining *k*-mer counts for mRNA and promoter regions, predictions were consistently more accurate generating separate feature sets for the promoter and mRNA regions (e.g., for 6bp *k*-mers resulting in 4^6^ or 4,096 * 2 = 8,192 features) that were both inputted into the model.

### Binary classification: Model explanation

Explainable AI was used to rank and select omic features as suggested by [75,76] and we investigated the explanations of the predictions for the DNA sequence-based ML model. Firstly, based on the best LightGBM model for the *Arabidopsis* Col-0 dataset from [8] on which the model was trained (i.e. 6,907+7,481= 14,388 transcripts and 8192 k-mer-based features). We applied the hyper-tuned LightGBM coupled up with an explainable AI algorithm, called Shapley Additive exPlanations - SHAP [38], as to predict and explain the class (circadian or non-circadian) of each transcript across the entire dataset. SHAP combines game theory with local explanation enabling accurate interpretations on why and how the model predicted a particular value (in our case a binary value) for a given instance. We used the python implementation of SHAP, version 0.35.0, available via the conda-forge channel (https://anaconda.org/conda-forge/shap). To obtain the appropriate SHAP explainer we combined shap.TreeExplainer with the hyper-tuned LightGBM model detailed in Table S2. Finally, we used the obtained SHAP explainer to compute SHAP values for the entire set of transcripts and *k*-mers. As we are performing a binary classification task, the SHAP explainer returned two SHAP values tables of the same dimension (number of transcripts x number of *k*-mer-based features), respectively for the class 0 (non-circadian) and the class 1 (circadian). In this manuscript we focus on the SHAP values for the class 1 – circadian. We used the SHAP summary plot function to produce Figure 3c that provides a global view of the local explanations when predicting class 1 (circadian) considering all samples in the dataset (training and test). Figure 3c shows the top 30 most impactful features/*k*-mers. Finally, we used the SHAP explainer to provides SHAP values, therefore explanations, for unseen transcripts from PHYA-E and the Col-0 and Ws-2 homologs of AT1G78040.3.

### Association of k-mers with TFBS, RNA binding motifs and miRNAs

We detail the closest matches of the *k-*mer’s to known TFBSs with p<0.05 as defined using Tomtom motif comparison with otherwise default settings [77]. The TFBSs used were *Arabidopsis* DAP-seq derived motifs [78]. We also detail the closest matches of the *k-*mer’s to known RNA binding motifs with p<0.05 as defined using Tomtom motif comparison with otherwise default settings [77]. Here, the RNA binding motifs used were from a systematic analysis of the RNA motifs recognized by RNA-binding proteins conducted by [79].The position of the *k-*mer is described as promoter or mRNA; if mRNA we express the percentage of mRNAs with the *k-*mer in their UTR region as a proportion of all genes with a UTR k-mer.

For each occurrence of the *k-*mer across the true positive circadian transcripts, 27nt sequence upstream and downstream of the 6-mer was extracted. These were then collated into a bwa-index for each 6-mer and 428 mature ath-miRNA sequences [80] were aligned to these indices using bwa-aln (maximum edit distance of 1, no gap opens or extensions allowed, seed sequence of 8nt, no edit distance permitted in seed sequence). SAM files were filtered to retain only those matches where the Arabidopsis query transcript was in the opposite orientation to the miRNA, included the k-mer and exhibited a maximum of 3 mismatches between putative target and miRNA. Candidate transcript/k-mer combinations were then tabulated with corresponding transcript information from Ensembl and miRNA annotation from miRBase [80].

### Filtering transcripts with most positive cumulative SHAP value

We summed the SHAP values individually for each transcript that our DNA sequence-based model accurately identified as circadian (true positives). The distribution of these cumulative SHAP values ranged from −0.27 to 9.61 with an average of 6.44 (Figure 4a). We filtered the circadian calls that were made with the most certainty according to the SHAP explanation (“most positive cumulative SHAP value”), removing those in the lower quartile Q1 i.e. those transcripts with a value lower than 6.29, leaving 5,536 of the transcripts where the most *k-*mers drive the prediction of circadian.

### Clustering genes using SHAP values

Clustering of genes based on SHAP values; for each gene we selected the top 5 most influential features or *k-*mers to its classification as circadian i.e., the 5 highest SHAP values. We then clustered the genes according to these profiles using hierarchical clustering with average linkage and Euclidean distance.

### Comparing morning/day/night/evening genes

We selected representative morning (phase 2-4hours), day (phase 9-11 hours), evening (phase 15-17 hours) and night (phase 21-23 hours) genes, by selecting those phases central to each of the groups as detailed since we define phase 0-6.99999 as morning, 7-12.999999 as day, 13-18.9999999 as evening, 19-24.999999 hours as night. For each group, across all genes we calculated the average SHAP value for each *k-*mer. We compared groups calculating the standard deviation between the groups for each *k-*mer. We ranked *k-*mers according to increasing variation between the four groups i.e. higher standard deviation or variability of *k-*mer importance.

### Identifying marker genes to tell the circadian time using a single transcriptomic timepoint

We developed a ML based pipeline to predict the circadian time (phase) at any single transcriptomic sampling timepoint using gene expression data from a set of marker genes. Here superior accuracy was achieved with an artificial neural network, it also allowed the simplistic implementation of a custom multioutput loss function, as such, we implemented this rather than the traditional ML methods used previously in this study. We note that this approach is more complex than our previous models, requiring custom code, as such we provide the code in a Jupyter Notebook and instructions to run this code at: https://github.com/JoshuaColmer/HallCircadian/. The three transcriptomic datasets used previously (Table S1) from [8], [9], and [10] were used for training, validation and testing respectively. The training dataset was used to learn the circadian time in hours, from the expression data from each transcriptomic timepoint individually, whilst the validation set was utilised for adjusting hyperparameters to reduce overfitting. The test set was used to estimate the error on unseen data for two different ecotypes Col-0 and Ws-2. Here, expression data was normalised by calculating transcripts per million (TPM) for increased uniformity between datasets, as in this experiment they were being directly compared. We removed genes whose expression distributions were too different between datasets based on the two-sample Kolmogorov-Smirnov test (q<0.05) and their minimum and maximum values, as well as removing low variance genes, using a threshold of 5, which was adjusted to minimise validation error. Since here, calculation of phase was critical to our predictions we extended our previous approach to quantify this more robustly; MetaCycle was used alongside cross-correlation with circadian time and autocorrelation, to quantify gene expression rhythmicity in the training dataset. The scores for each metric were combined using a Gaussian copula yielding one score per gene. The top-ranking *n* genes were taken forward for model training and further feature selection where *n* was adjusted to minimise validation error. As previously MinMaxScaler was used to scale the features from 0 to 1, fitted on the training set and applied to validation and test sets.

We created a shallow neural network using TensorFlow (v2.0.0) [81] comprising three fully connected layers with ReLU activation functions and 32, 128, 512 and 2 neurons respectively followed by a 2 neuron softmax layer. The learning rate, number of training epochs and architecture of the network were optimised using the hyperas (v0.4.1) package to minimise the loss for the validation dataset. Due to the cyclical nature of the target (time 0-24 hours), standard regression loss functions were not suitable for this task. To quantify the error in the predictions, we defined the loss function as the squared angle between actual circadian time and predicted circadian time after being transformed onto a unit circle.

We used feature selection as previously to select *n* circadian genes for model training, prioritizing weighted representation of genes from each of the 8 expression sub-clusters generated by the WGCNA gene co-expression network analysis [54]. We hoped this would improve generalisation and robustness of the model as the similarity between features would be reduced and the diversity of features should enable the neural network to engineer more complex representations of the expression data compared to if all features belonged to the same phase cluster.

## Declarations

### Ethics approval and consent to participate

Not Applicable

### Consent for publication

Not Applicable

### Availability of data and materials

All of the datasets used in this analysis are detailed in Table S1. All previously published datasets have details for data generation in the relevant associated publication. For the wheat time course: reads are available from the NCBI Sequence Read archive under project name PRJEB40948 at https://www.ebi.ac.uk/ena/browser/view/PRJEB40948. The algorithms and hyperparameters used for the detailed ML models are stated in Table S2.

### Code availability

For our circadian time prediction custom code is required, as such we provide the code in a Jupyter Notebook and instructions to run this code at: https://github.com/JoshuaColmer/HallCircadian/.

### Authors’ contributions

This work was conceived by LJG, AH, RK with additional ideation from EPK. LJG led the analyses with assistance from RRP (bioinformatics analyses), JMC (timepoint reduction and *k*-mer spectra ML), JC (clock function ML), HR (investigation and interpretation of results) and APC (ML guidance and SHAP analysis). Laboratory work for the wheat dataset was carried out by SD with assistance from HR. LJG wrote the manuscript with guidance from AH and HR, a section written by JC plus methods sections written by RRP and APC and general assistance from all other authors. All authors read and approved the final manuscript.

## Acknowledgements

Not Applicable

## Supplementary Information

Supplementary Notes 1-6

Supplementary Figures 1-7

Supplementary Tables 1-15

Supplementary References 1-36

