## Supplementary Information for "Interpreting machine learning models to investigate circadian regulation and facilitate exploration of clock function"

Supplementary Notes 1-6

Supplementary Figures 1-7

Supplementary Tables 1-15

Supplementary References 1-36

**Supplementary Note 1: Reasoning behind MetaCycle labelling of transcripts and cut-offs**

We use MetaCycle as a "gold-standard" circadian/non-circadian labeller for our transcripts to allow us to quantify the accuracy of transcript classification downstream by our ML models (Wu et al., 2016). As such, we need to be sure of the accuracy of MetaCycles labelled transcripts. MetaCycle is known to be effective with densely sampled timepoints like the 12 we use here (or more). However, the accuracy of circadian gene identification between software's is widely debated and a detailed analysis of software accuracy on densely sampled time-series is beyond the scope of this work-with our focus on reducing the need for transcriptomics. As such, partly to bypass this issue, for model training to label transcripts we used the 7,734 most high confidence circadian transcripts defined by MetaCycle using 12 timepoints. We balanced stringency of thresholds with having sufficient training examples for ML. As such, we denoted those transcripts as most high confidence with a q-value of <0.02 rendering them highly likely to be circadian (standard recommended cut-off q<0.05).

In support of this, we compared transcript rhythmicity detection between MetaCycle and another software BooteJTK that, although not so widely cited, as of yet, is reported to give significantly higher accuracy than MetaCycle (Hutchinson et al., 2018). For this comparison we used parameters to run BooteJTK that most closely represented those used for MetaCycle and the exact same input data. We observe that 98% of our high confidence circadian transcripts that were identified by MetaCycle were also defined as such by BooteJTK. This highlights that using a stringent cut-off to define our high confidence circadian transcript set minimises the impact of software choice to label our transcripts.

**Supplementary Note 2: ML classification strategy**

We trained ML classifiers to predict if a transcript was circadian or non-circadian in a binary classification system using the extremes of circadian (rhythmic) and non-circadian (non-rhythmic) transcripts as defined by using specific q-value thresholds to filter the MetaCycle output. As previously mentioned, firstly, this largely ensured that true positives and true negatives were used for model training independent of our choice of software for labelling (Supplementary Note 1). However, we do lose information from the models by training the transcriptomic model on 15,468 (7,734*2) out of a possible 44,963 transcripts. Critically, although we have employed this tactic for model training, we did so with the anticipation that training on strong examples would not negate our potential to also classify cases in the *grey area* in the middle. In support of this, when we apply our trained ML model (12 transcriptomic timepoints) on all 44,963 transcripts we still show a classification accuracy of 90.3% overall (87% circadian class, 92% non-circadian class). Furthermore, if we focus on the 36,217 or 81% of transcripts that MetaCycle can also confidently quantify, our trained transcriptomic ML model shows an accuracy of 94% on the circadian class and 95% on the non-circadian class. We can conclude that training on the extremes of rhythmic and non-rhythmic translates effectively to the other intermediate transcripts.

**Supplementary Note 3: Comparing transcript-specific explanation for PHYA-E classifications**

We compared the SHAP impact values between each of the PHY transcripts A/B/D/E (circadian) and PHYC (non-circadian). Here a high comparative number translates to regulatory elements being more impactful in predicting PHYA/B/D/E to be circadian but also typically more impactful in predicting PHYC to be non-circadian. Therefore, the change in frequency of these regulatory elements is closely associated with, and more likely to be responsible for, the circadian/non-circadian predictive differences according to our model. Selecting the top three highest values per comparison (PHYA versus PHYC, PHYB versus PHYC, PHYD versus PHYC and PHYE versus PHYC) encompassed the outliers with the largest differences in SHAP values shown in Figure S3a. We found that one *k*-mer was significantly different in all four comparisons (GGTAGA) and three others were in a minimum of two comparisons (CCGTCG, TTTC[A/T]G and TCTCCG) (Table S9). GGTAGA and TTTC[A/T]G had the lowest SHAP values recorded for PHYC (driving its prediction of non-circadian) and both of their frequencies of occurrence in the mRNA correlate with the prediction probability of the circadian class across the transcripts (>0.3) (Figure S3b). GGTAGA was present only in the exonic regions of PHYC but also in PHYD (Figure 5) and both showed the lowest predicted probabilities of being circadian class (0.38 and 0.55 respectively). TTTCTG sites were found within the PHYC 5'UTR and just after the TSS but were missing from these regions in the other transcripts (again except for one occurrence in the 5'UTR of PHYD). Conversely, AAATAA was present in the 5'UTR of PHYA and PHYE but missing from PHYC and TCTCCG was present in the 5'UTR of PHYB but missing from PHYC.

**Supplementary Note 4: Labelling of Ws-2 transcripts as circadian/non-circadian**

No confident genome-wide circadian/non-circadian labelling of Ws-2 genes currently exists for us to measure our predictive accuracy. Graf et al. (2017) generated transcriptomic information for both Ws-2 and Col-0 across two timepoints (0 and 12 hours after constant light) (Table S1). As such, we used their Col-0 time-series data, combined with our gold-standard Col-0 circadian/non-circadian target labels determined previously with 12 timepoints, to train a model that we could use downstream to label Ws-2. We trained a series of ML classifiers using the Col-0 labelled transcriptomic data using the same transcripts used previously in our ML models (see Methods). The best model was generated with LightGBM showing (using best parameters after fine tuning): F1 scores of 0.738/0.777 for non-circadian/circadian respectively on the training data, F1 scores of 0.724/0.760 on the (held out) test data and a mean F1 cross validation score of 0.750 on the test data.

Although we achieved relatively high accuracy predicting Col-0 transcript labels from such a low number of transcriptomic timepoints, this model was built to label Ws-2 using its two transcriptomic timepoints and these Ws-2 labels were needed to measure the accuracy of our DNA sequence-based model predictions on Ws-2. As such, accurately labelled Ws-2 is a priority to provide a precise measure of our DNA sequence-based model's predictions. To identify the models most confident transcript predictions, for this case of binary classification, we calculate two probabilities for a given prediction, the probability of the label being class 0-non-circadian and of it being class 1-circadian. These predictive probabilities sum up to 1 across the classes and we translate them into a percentage. We determined that higher prediction probabilities for datapoints (for one of the two classes) associated with higher classification accuracy and that applying a threshold to predictions (prediction probability of 80% or more in one class) allowed our model to achieve a more acceptable >90% accuracy on predictions for 6,866 Col-0 transcripts (Figure S4). We next applied this Col-0 ML transcriptome-based model, using our prediction probability threshold, to the transcriptomic data from Ws-2 to label Ws-2 transcripts. Since correct labelling of Ws-2 transcripts is unknown, we have no way to measure accuracy but checked for agreement of circadian/non-circadian classifications between Ws-2 and Col-0, observing agreement for 84.6% of transcripts. Given that the average accuracy of the ML model being used was ~90%, the number of transcripts with conserved classifications between Col-0 and Ws-2 is in line with what we may expect from ecotypes.

Having defined circadian/non-circadian labels for the Ws-2 transcripts, we used these as test datapoints for our DNA-sequence based ML model (trained on Col-0), alongside *k*-mer profiles that we generated *de-novo* for the mRNA and promoter sequences associated with each Ws-2 transcript, as per previous methods, with additional integration of Ws-2 SNP information to evolve the Col-0 reference (see Methods).

**Supplementary Note 5: Performance of DNA sequence-based model on Ws-2**

Returning to our case study of 41 known circadian genes, our DNA sequence-based ML model produced a high accuracy of 82.5% on their Ws-2, homologs which is only ~10% lower than with Col-0 (Table S11). We considered that some Ws-2 homologs of Col-0 known circadian genes may not be rhythmic so this accuracy could be an underestimate. However, this analysis highlights how our model generalizes to new ecotypes since although, using Col-0, a proportion of the 41 genes (75.6%) were used in model training, no Ws-2 information was incorporated (only Ws-2/Col-0 homologs will be present), therefore these Ws-2 transcripts are all unseen datapoints to the model. Of the 10 of the 41 genes where even no Col-0 homolog was used for model training; we were still able to achieve accuracy up to 83.3% on Ws-2

Previously, as a measure of the worst-case scenario for predictions, we focused on 10 of the 41 known circadian genes from the literature that were not used to train either of our ML models, largely due to them being problematic (too low amplitude) for our first step of labelling or classification by MetaCycle (q>0.01), even using 12 timepoints. For the Ws-2 sequences corresponding to these 10 genes that are already known to be problematic to classify, the model will have never seen even the Col-0 homolog of the gene, and, as such, these will be much more difficult to classify for Ws-2; we recorded an accuracy of 66.7% which is >20% lower than with Col-0 (Table S11). Notably, our 66.7% accuracy is still significantly higher than that obtained on these genes using MetaCycle (20.0%) and ML (50.0%) on the Ws-2 transcriptomic data directly. We applied an 80% predictive probability threshold to the problematic 10 genes, only considering those predictions where our DNA sequence-based model was highly confident (defined in Supplementary Note 4). Our model was able to confidently classify 67% of these most challenging unseen transcripts for Ws-2 and for this confident group our accuracy rate was increased to 83.3%, <7% lower than we saw with Col-0. The consideration of predictive probability allows us to label the majority of transcripts while also providing a confidence measure in the prediction to ensure that a user can trust the predictions.

**Supplementary Note 6: Top ranked k-mers differentiating Ws-2 and Col-0 RPP7**

The top 5 ranked *k*-mers that were most different between Ws-2 and Col-0 (according to SHAP impact), and were ultimately used to evolve the Ws-2 homolog, were all found in the mRNA and we identified the closest matching annotations for each of them considering both TFBS and miRNA binding sites. The top 5 k-mers and their matches were as follows: TGATCA.1 matched miR5633 (p=0.009) that targets AT1G12280, a NB-LRR disease resistance protein SUPPRESSOR OF MKK1 MKK2 2 (SUMM2); GGTAGA.1 matched the MYB61 transcription factor (p=0.003) that regulates light-induced stomatal opening and dark-induced stomatal closure that can be utilized as a method of plant defence (Nagatoshi et al., 2016); GAATGT.1 matches another as-of-yet undefined MYB family transcription factor AT2G40260 (p=0.001) known to interact with disease resistance associated Mitogen-activated protein kinase 5 (MAPK5); TAAGAT.1 matched TARGET OF EAT 1/2 (TOE1/2) (p=0.005), TOEs counteract the promotion of flowering by CO (Zhang et al., 2015); finally, TTTCAG.1 matched AT1G63040 (AtERF026) (p=0.003) that has been linked to response to downy mildew pathogen response (Huibers et al., 2009).

**Supplementary Note References:**

Hutchison AL, Allada R, Dinner AR. Bootstrapping and Empirical Bayes Methods Improve Rhythm Detection in Sparsely Sampled Data. J Biol Rhythms. 33(4):339-349 (2018)

Wu, G., Anafi, R. C., Hughes, M. E., Kornacker, K., & Hogenesch, J. B. MetaCycle: an integrated R package to evaluate periodicity in large scale data. *Bioinformatics*, **32(21)**, 3351–3353. (2016)

Graf, A., Coman, D., Uhrig, R. G., Walsh, S., Flis, A., Stitt, M., & Gruissem, W. Parallel analysis of Arabidopsis circadian clock mutants reveals different scales of transcriptome and proteome regulation. *Open biology*, **7(3)**, 160333. (2017)

Nagatoshi, Y., Mitsuda, N., Hayashi, M., et al. GOLDEN 2-LIKE transcription factors for chloroplast development affect ozone tolerance through the regulation of stomatal movement. *Proc Natl Acad Sci U S A*. **113(15),** 4218-4223. (2016)

Zhang, B., Wang, L., Zeng, L., Zhang, C., Ma, H. Arabidopsis TOE proteins convey a photoperiodic signal to antagonize CONSTANS and regulate flowering time. *Genes Dev.* **29(9),** 975-987. (2015)

Huibers, R.P, de Jong, M., Dekter, R.W., Van den Ackerveken, G. Disease-specific expression of host genes during downy mildew infection of Arabidopsis. *Mol Plant Microbe Interact*. **22(9)**, 1104-1115. (2009)

### Supplementary Figures

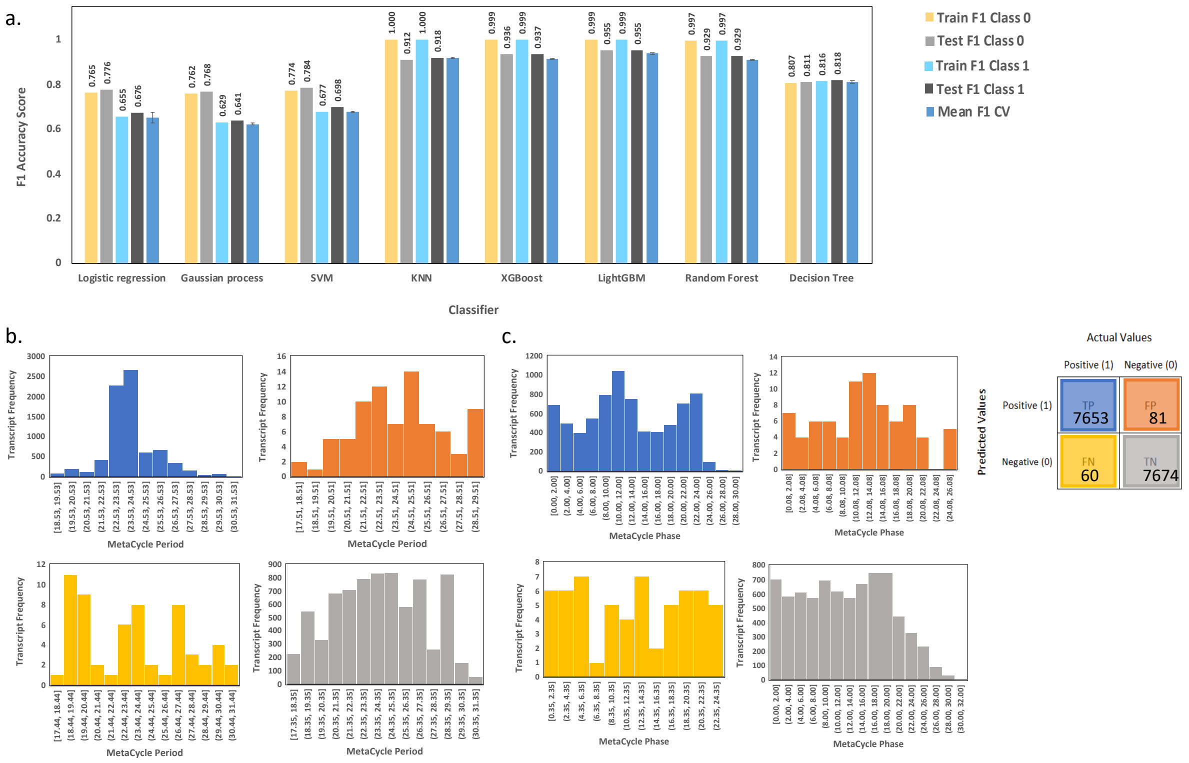


**Figure S1. Results from Arabidopsis circadian/non-circadian best ML binary classification analysis with 12 transcriptomic timepoints. (a)** Bar charts showing the F1 accuracy scores for the training and test datasets (using best parameters) and the mean F1 accuracy scores on the test data after 5-fold cross validation with the standard deviation shown as error bars. Labels for different bar colours are detailed in legend to the right of the plot, Class 0=Non-circadian and Class 1=Circadian. Histograms in **(b-c)** all relate to the best ML model generated wiith LightGBM from 12 Arabidopsis transcriptomic timepoitns. The histograms are colour coded as per the confusion matrix shown in the legend above i.e. showing where our model assigned True Positive labels (TP), False Positive labels (FP), False Negative labels (FN) and True Negative labels (TN). The histograms show the frequency of transcripts that had various **(b)** period lengths or **(c)** phases assigned to them by Metacycle.

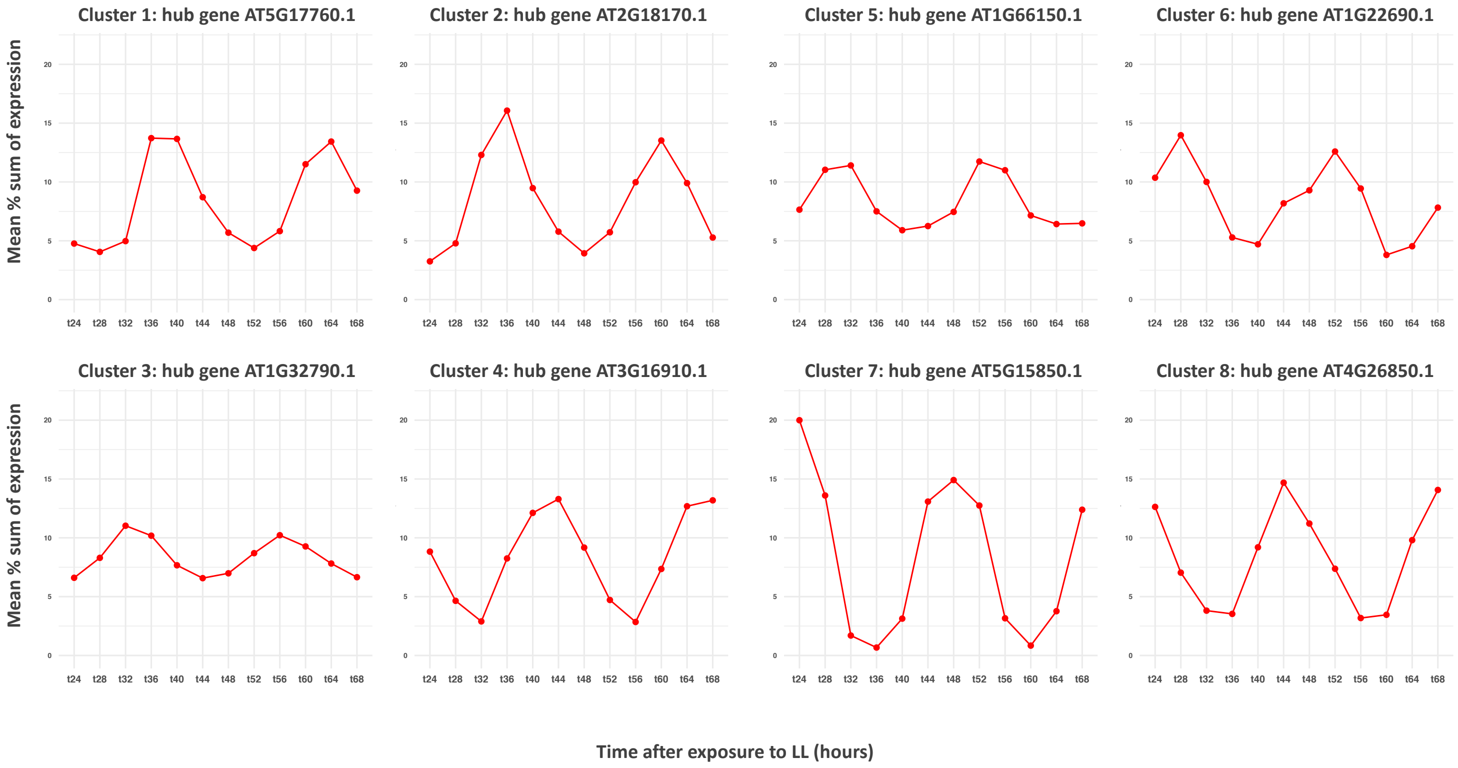


**Figure S2. Clustering Arabidopsis (Col-0) circadian transcripts according to transcriptomic profiles.** Weighted gene co-expression network analysis (WGCNA) was used to divide the circadian transcripts into 8 different modules in scale-free networks, where the transcripts in each module have the same expression pattern. Hub genes with high connectivity in the module were identified and the transcriptomic profiles of these genes for each of the 8 modules are shown here.

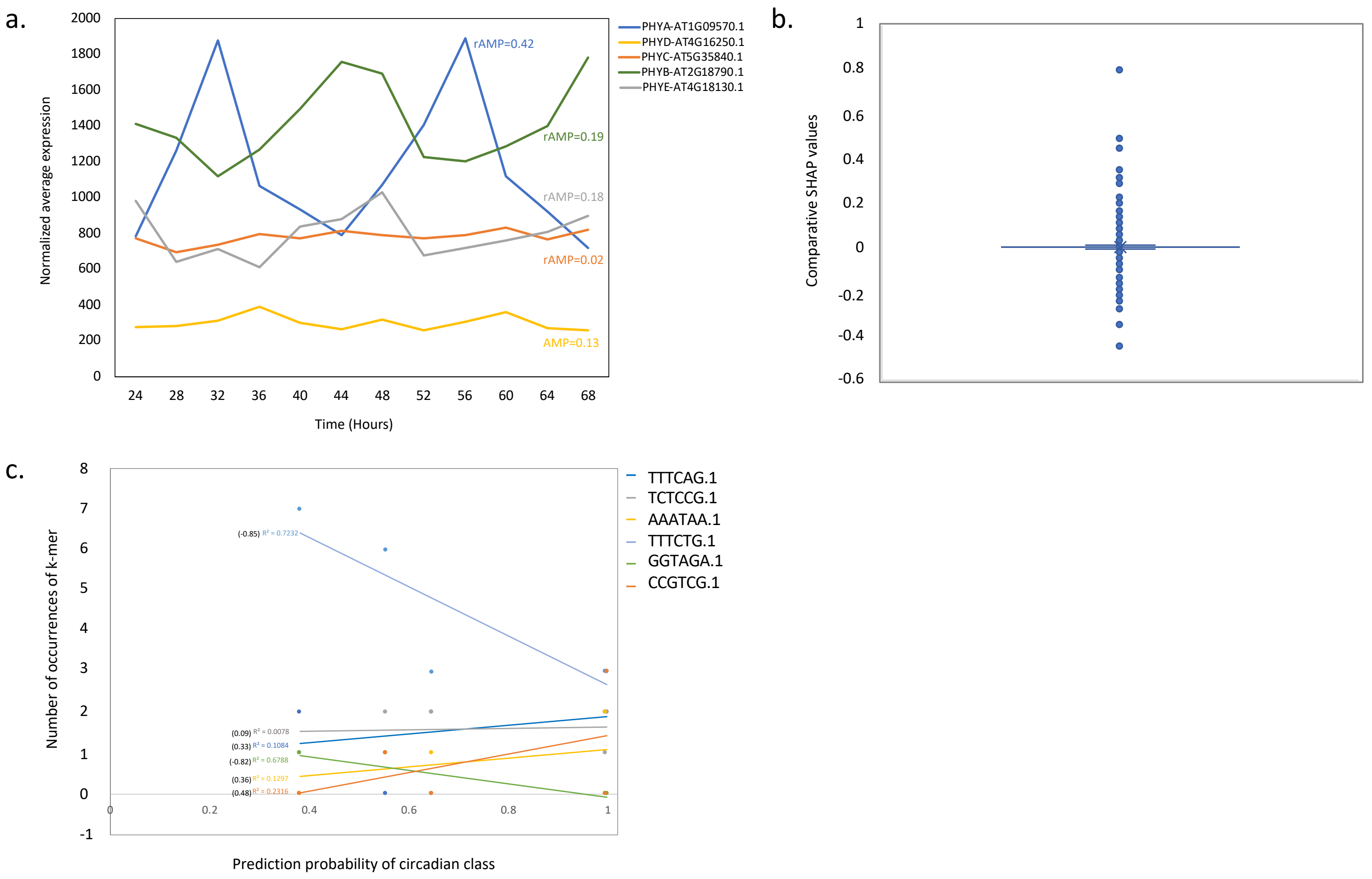


**Figure S3. Results from SHAP explanation of Arabidopsis (Col-0) circadian/non-circadian best ML binary classification DNA sequence-based analysis. (a)** Line plots of the normalized expression (averaged across the replicates) for the transcripts corresponding to PHYA-E with respect to time of sampling. Relative amplitudes of the expression profiles (as calculated by MetaCycle) are shown in brackets. **(b)** box plot showing the range of computed values when we compared the SHAP explanations between each of the PHY primary transcripts A/B/D/E and PHYC i.e. PHYA versus PHYC, PHYB versus PHYC, PHYD versus PHYC and PHYD versus PHYC. (**c**) xy scatter plot of the frequency of occurrence of each k-mer in the mRNA versus the prediction probability of the circadian class associated with that *k*-mer. Lines shown are linear trend lines plus r2 values and Pearson correlation coefficients shown in brackets.

a.

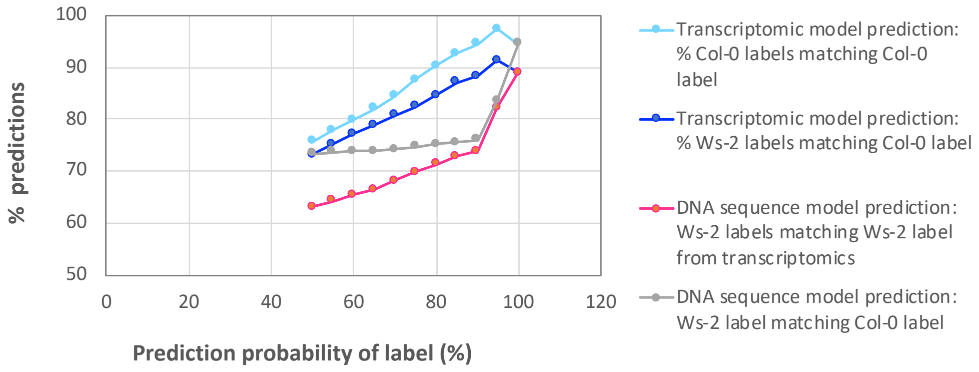


b.

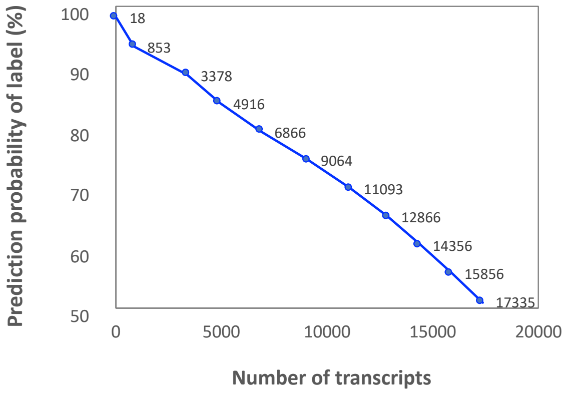


**Figure S4. Results from Arabidopsis (Ws-2/Col-0) circadian/non-circadian best ML binary classification analysis.** Using the dataset generated by Graf et al. (2017) that had two transcriptomic timepoints for Col-0 and Ws-2. **(a)** x-y scatter showing the prediction probability of the label assigned to each transcript on y-axis versus the percentage of predictions that fitted each of the following categories (see legend) on the x-axis; **(light blue/dark blue**) Application of "Transcriptomic model" that was trained with two timepoints from Col-0 using the original gold-standard labels defined previously using 12 timepoints. **(light blue**) % of predicted labels for Col-0 matching known labels. **(dark blue)** % of predicted labels for Ws-2 matching known labels for Col-0. **(raspberry/grey**) Application of "DNA sequence model" that was trained with k-mer profiles from Col-0 using the original gold-standard labels defined previously using 12 timepoints. **(raspberry)** % of predicted labels for Ws-2 matching known Ws-2 labels (defined by transcriptomic model). **(grey)** % of predicted labels for Ws-2 matching known Col-0 labels. **(b)** For increasing prediction probability (%) of the label as seen in **(a)** here we show the number of transcripts that had labels predicted at a minimum probability of each threshold from 50-100% (highest class), independently of whether the prediction was accurate or not.

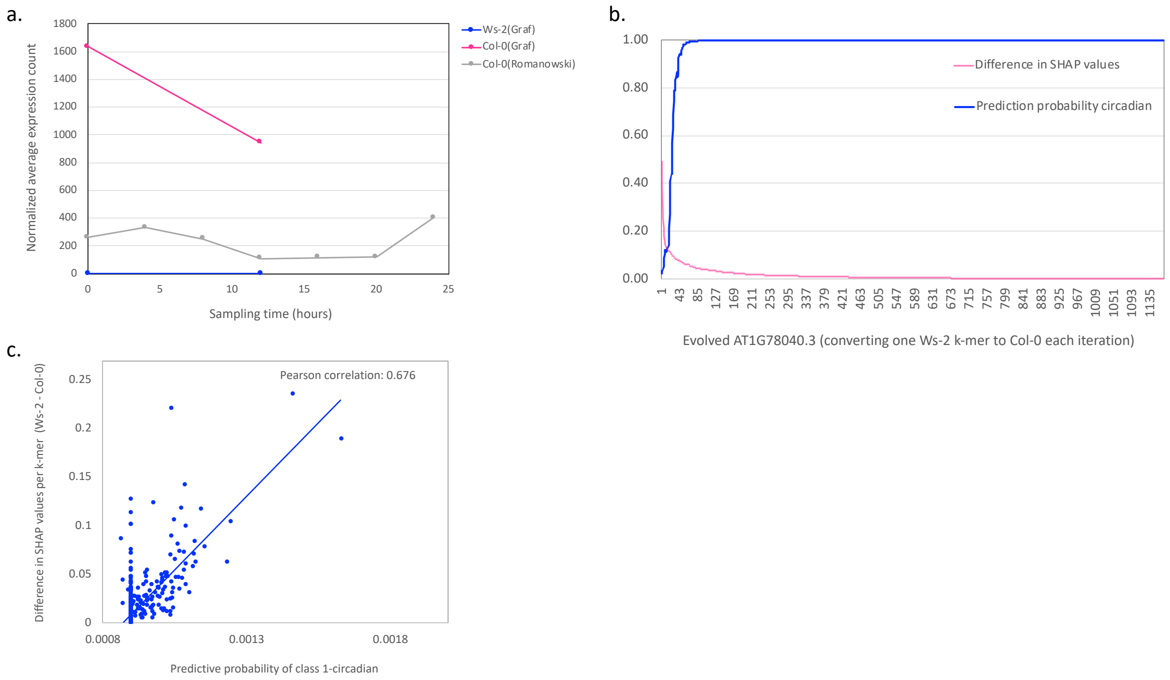


**Figure S5. Comparing transcript AT1G58602.1 from Arabidopsis (Col-0/Ws-2) circadian/non-circadian DNA sequence-based ML binary classification analysis.** Here we focus on the transcript AT1G58602.1 that was predicted circadian by the transcriptomic information and *k*-mer based model (both predictive probability >0.8) for Col-0 but predicted non-circadian in Ws-2. **(a)** Showing the transcriptomic profiles available for this trancript in Ws-2 and Col-0 and their sources. **(b)** We sequentially evolved the *k*-mer spectrum for AT1G58602.1 in Ws-2 a k-mer at a time to match Col-0 more and more with each iteration. Only *k*-mers where the SHAP explanation showed a positive impact for a circadian call in Col-0 and a negative impact for a circadian call in Ws-2 were used and *k*-mers were ordered with the largest size difference between SHAP values in Col-0 and Ws-2 first. For each evolved transcript we re-ran it through the DNA sequence-based classification model and here we show the predictive probability of the circadian class for each evolved transcript. We also show the corresponding difference between SHAP values in Col-0 and Ws-2 for each *k*-mer. **(b)** We plot the predictive probability of the circadian class for each evolved transcript versus the difference between SHAP values in Col-0 and Ws-2. Here, conversely to in **(b)** each evolved transcript has one *k*-mer converted from Ws-2 to Col-0 (rather than sequentially accumulating changes) so the number of evolved transcripts matches the number of *k*-mers where the SHAP explanation showed a positive impact for a circadian call in Col-0 and a negative impact for a circadian call in Ws-2 (1168).

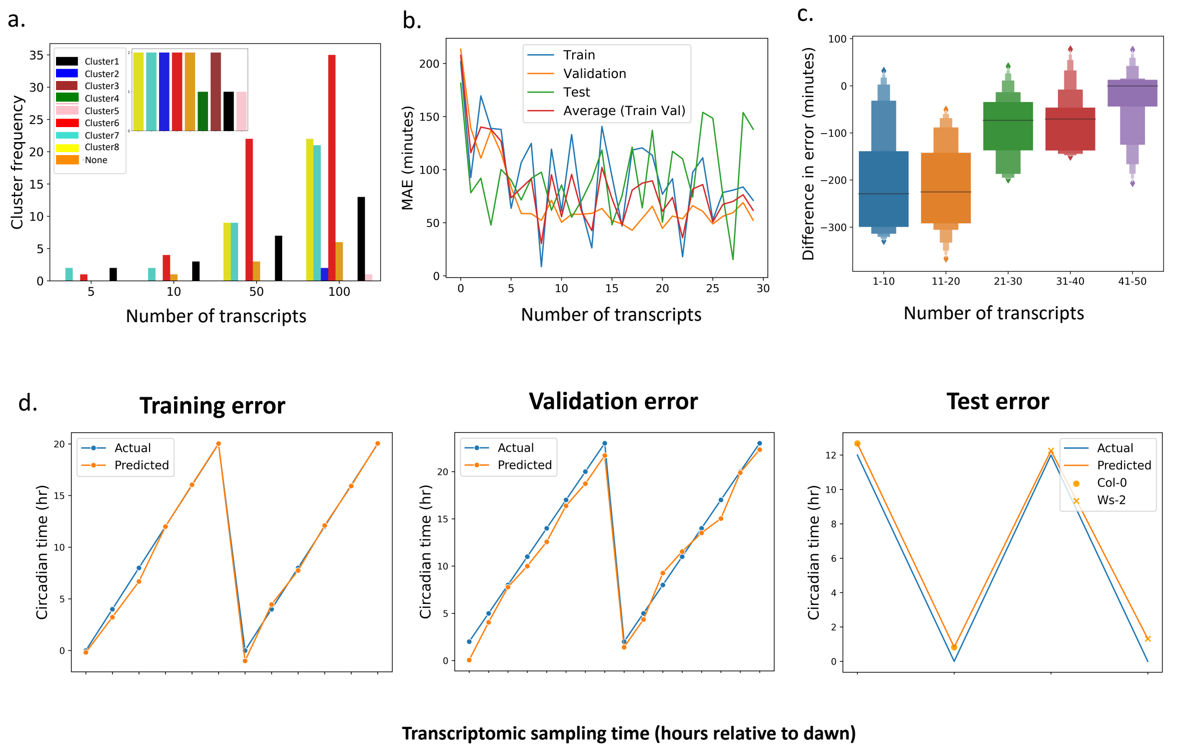


**Figure S6. Identifying marker genes that tell the time using a single transcriptomic timepoint. (a)** Bar chart to show the frequency distribution of the circadian transcripts for various sized subsets according to their presence in the 8 expression sub-clusters generated by the WGCNA gene co-expression network analysis. Inset is the same bar chart for the final 15 selected transcripts after the cluster-based sequential feature selection i.e. to even out cluster representation. **(b)** line plot displaying the mean absolute error (MAE) predictive results of circadian time after the cluster-based sequential feature selection which shows the error for each dataset at each iteration (right). **(c)** A letter value boxplot displaying the distribution of improvements made by using the cluster-based sequential feature selection on the test set for different numbers of transcripts, negative scores indicate it reduced the error, positive scores indicate it increased the error. **(d)** Three line plots for training, validation and test predictions compared to ground truth data using our final subset of 15 transcripts and optimized hyperparameters.

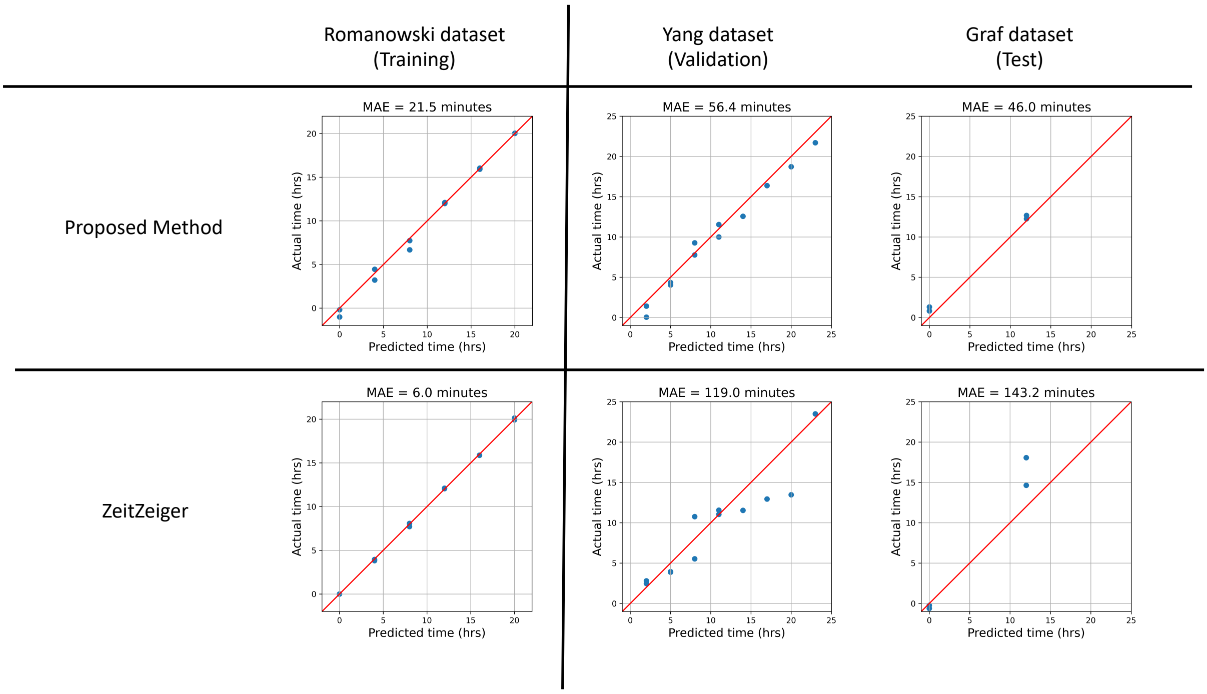


**Figure S7. Comparing our proposed method for telling the circadian time to the state-of-the-art.** Six scatterplots displaying the predictive performance of our proposed method compared to the performance of ZeitZeiger on the same three datasets. Each blue point is a sample and the red lines represent where the actual time is equal to the predicted time. The title for each scatterplot is the mean absolute error (MAE) of the predictions of the given model and dataset combination, where the MAE is the average distance between the actual time and the predicted time in minutes.

### Supplementary Tables

**Table S1. Datasets used in this analysis.** Showing the data sources and features.

| SOURCE | ECOTYPE | MATERIAL | BIOLOGICAL REPLICATES (PER TIME POINT) | DELAY IN SAMPLING AFTER TRANSFER TO LL (HRS) | TIMEPOINT NUMBER | SAMPLING FREQUENCY (HRS) | DATA TYPE |
|---|---|---|---|---|---|---|---|
| Romanowski et al. (2020) | Col-0 | Areal | 2 | 24 | 12 | 4 | RNA-seq 361 million Illumina reads |
| Yang et al. (2020) | Col-0 | Seedlings | 2 | 2 | 8 | 3 | RNA-seq Illumina reads |
| Graf et al. (2017) | Col-0 and Ws-2 | Whole rosettes | 2-3 (9 plants per replicate) | 0 | 2 | 12 | RNA-seq 20-50 million Illumina reads (per replicate) |
| Rees et al. (2020) | Wheat-Cadenza | Leaf-seedlings | 4 | 72 | 24 | 2 | RNA-seq 60 million Illumina reads (per replicate) |

**Table S2. Best performing classification models from ML analyses to predict if a transcript is circadian or non-circadian.** Detailing the parameter sets used for our best performing models.

| FEATURE SET | CLASSIFIER | HYPERPARAMETERS |
|---|---|---|
| **MODEL 1**<br><br>**12 Arabidopsis timepoints** | LightGBM | random_state=42, boosting_type='gbdt', class_weight=None, colsample_bytree=1.0, importance_type='split', learning_rate=0.6554916745454337, max_depth=7, min_child_samples=50, min_child_weight=0.001, reg_lambda=0.0, silent=True, subsample=0.7, subsample_for_bin=200000, subsample_freq=0, num_leaves=20, n_estimators=450, min_data_in_leaf=50, min_split_gain=0.0, objective=None, reg_alpha=0.0, verbose=False |
| **MODEL 2**<br><br>**6bp *k*-mer profile promoter+cDNA** | LightGBM | boosting_type='gbdt', class_weight=None, colsample_bytree=1.0, importance_type='split', learning_rate=0.13853569656385206, max_depth=9, min_child_samples=25, min_child_weight=0.001, min_split_gain=0.0, n_estimators=350, num_leaves=100, objective=None, random_state=42, reg_lambda=0.0, silent=True, subsample=0.5, subsample_for_bin=200000, subsample_freq=0 |

**Table S3. Arabidopsis circadian/non-circadian comparative ML binary classification analysis to reduce the number of transcriptomic timepoints.** For our best ML model, we reduced the number of timepoints sequentially from 12 down to 3. To obtain each reduced set of timepoints, we compare using chi-square (Chi2) and eli5 (Eli5) feature selection with every possible random feature combination (Random). Here we show the best F1 accuracy score on the test data after 5-fold cross validation for each set of reduced timepoints.

| Number of time points | Model used for classification | Feature Selection method | Mean F1 Accuracy on CV | SD of F1 accuracy on CV | Feature Selection method | Mean F1 Accuracy on CV | SD of F1 accuracy on CV |
|---|---|---|---|---|---|---|---|
| 12 | LightGBM | NA | 0.939 | 0.003 | NA | 0.939 | 0.003 |
| 11 | LightGBM | CHI2 | 0.936 | 0.002 | Eli5 | 0.938 | 0.003 |
| 10 | LightGBM | CHI2 | 0.919 | 0.002 | Eli5 | 0.932 | 0.003 |
| 9 | LightGBM | CHI2 | 0.907 | 0.005 | Eli5 | 0.911 | 0.004 |
| 8 | LightGBM | CHI2 | 0.867 | 0.005 | Eli5 | 0.894 | 0.002 |
| 7 | LightGBM | CHI2 | 0.856 | 0.005 | Eli5 | 0.860 | 0.003 |
| 6 | LightGBM | CHI2 | 0.823 | 0.004 | Eli5 | 0.836 | 0.007 |
| 5 | LightGBM | CHI2 | 0.799 | 0.005 | Eli5 | 0.809 | 0.005 |
| 4 | LightGBM | CHI2 | 0.781 | 0.005 | Eli5 | 0.767 | 0.007 |
| 3 | LightGBM | CHI2 | 0.753 | 0.007 | Eli5 | 0.755 | 0.005 |

| Number of time points | Feature Selection method | Mean F1 Accuracy on CV | SD of F1 accuracy on CV | Test F1 Accuracy circadian class | Test F1 Accuracy non-circadian class |
|---|---|---|---|---|---|
| 12 | NA | 0.939 | 0.003 | 0.944 | 0.944 |
| 11 | Random | 0.934 | 0.001 | 0.938 | 0.939 |
| 10 | Random | 0.929 | 0.002 | 0.928 | 0.929 |
| 9 | Random | 0.922 | 0.003 | 0.925 | 0.925 |
| 8 | Random | 0.915 | 0.005 | 0.912 | 0.913 |
| 7 | Random | 0.901 | 0.002 | 0.897 | 0.900 |
| 6 | Random | 0.886 | 0.003 | 0.878 | 0.878 |
| 5 | Random | 0.856 | 0.005 | 0.846 | 0.849 |
| 4 | Random | 0.815 | 0.003 | 0.801 | 0.805 |
| 3 | Random | 0.792 | 0.004 | 0.776 | 0.778 |

**Table S4. Arabidopsis circadian/non-circadian comparative analysis to reduce the number of transcriptomic timepoints.** We reduced the number of timepoints using MetaCycle from 12 down to 6 in several different ways as detailed. For 3 timepoints we compare our models to the state-of-the-art method from Spörl et al. (2012). Here we show the comparative definition of circadian transcripts for each set of reduced timepoints compare to the analysis with the original 12 timepoints.

| **Number of time points** | **Model used to classify** | **Feature Selection method** | **Number of circadian transcripts (q<0.05)** | **% of 9394 Circadian transcripts identified using 12 timepoints if q<0.05** | **% of 9394 Circadian transcripts identified using 12 timepoints if p<0.05** | **Mean F1 accuracy score on CV for ML model (%)** |
|---|---|---|---|---|---|---|
| 12 | MetaCycle | NA | 9394 | 100 | 100 | 93.9 |
| 6 | MetaCycle | 4hourly/day1 | 522 | 4.9 | 58.2 | 88.4 |
| 6 | MetaCycle | 4hourly/day2 | 132 | 1.3 | 52.5 | 88.3 |
| 6 | MetaCycle | 8hourly/2days (24-64h) | 712 | 4.0 | 57.1 | 83.9 |
| 6 | MetaCycle | 8hourly/2days (28-68h) | 2455 | 14.8 | 63.7 | 83.9 |
| 6 | MetaCycle | 4hourly/1day (36-56h) | 101 | 0.9 | 50.0 | 88.6 |

| **Number of time points** | **Method used to classify** | **Feature Selection*** | **Number of simulated biological replicates** | **Standard deviation between biological replicates** | **Accuracy (AUC)** | **Test set AUC score for ML model ****** |
|---|---|---|---|---|---|---|
| 3 | Spörl et al. | 9.30/14.30/19.30 | 20 | 1 | 0.87 | 0.78 |
| 3 | Spörl et al. | 9.30/14.30/19.30 | 10 | 1 | 0.82 | 0.78 |
| 3 | Spörl et al. | 9.30/14.30/19.30 | 2 | 1 | 0.78** | 0.78 |
| 3 | Spörl et al. | 9.30/14.30/19.30 | 2 | 1.64 | 0.66*** | 0.78 |

* Timepoint matching not possible so closest equivalents selected.
** Extrapolated AUC: to align the number of biological replicates with our Romanowski ML model training set
*** Extrapolated AUC: to align with standard deviation between biological replicates observed with our Romanowski ML model training set
**** Here the AUC score was computed for our best ML model with 3 time points to allow accurate comparison with Spörl et al.

**Table S5. The top 30 most impactful *k*-mers for our best Arabidopsis circadian/non-circadian binary classification ML model (LightGBM).** The top 30 most impactful *k*-mers for predicting class 1 (circadian) considering all samples in the dataset as calculated using SHAP. A *k*-mer is coloured red if higher frequencies of it contribute to a circadian classification, the *k*-mer is coloured blue if lower frequencies of it contribute to the circadian classification of a transcript and the *k*-mer is coloured black if the direction of contribution is unclear. We detail the closest matches of the *k*-mer's to known Arabidopsis TFBS (p<0.05) (see Methods). We describe the TF that binds to the TFBSs. The position of the *k*-mer is described as promoter or mRNA; if mRNA we express the percentage of all tested mRNAs with the *k*-mer in their UTR region. Note that these are the top 30 most impactful *k*-mers for predicting circadian transcripts, this does not mean necessarily that all circadian transcripts will have that *k*-mer and that it will be critical for its rhythmicity-only that overall they are impactful across more transcripts than other *k*-mers. For the two *k*-mers with miRNA targets, we report the associated miRNA's as determined using the stated alignment methods and thresholds in the main text and methods section; we also report the references that directly link the miRNA's to circadian rhythmicity or regulation. For *k*-mers matching RNA binding motifs (RNA-BMs) recognized by RNA-binding proteins, we report the associated RNA-binding proteins as determined using the stated alignment methods and thresholds in the main text and methods section; we also report the references that directly link the RNA-binding proteins to circadian rhythmicity or regulation.

| Rank | K-mer | Position | Associated TFBS Motif Sequence | Associated TFBS Motif P-value | Associated TFBS Motif E-value | Description of TFs binding associated TFBS motif<br>Description of miRNA targets of *k*-mer if appropriate |
|---|---|---|---|---|---|---|
| 1 | TCTCCG.1 | mRNA UTR:28.18% | AAATCTCCGGCGACG | 0.0002 | 0.184 | Trihelix_tnt.AT3G58630_col_a_m1 and Trihelix_tnt.AT5G05550_colamp_a_m1: AT3G58630 has a protein-protein interaction[1] with LIGHT INSENSITIVE PERIOD1, LIP1 which influences the light input pathway of the plant circadian network[2]. BioGRID[3]: AT5G05550 interacts with Circadian related TCP2[4], CONSTANS-LIKE 11[5] and RVE2[6] |
| 2 | TCTTCT.1 | mRNA UTR:93.34% | GGAATCTTCTA | 0.002 | 1.69 | G2like_tnt.At1g25550_col_m1:<br>Myb-like transcription factor family protein[7] |
| 3 | CCGTCG.1 | mRNA UTR:11.23% | CCGTCGAATTCACGGCA<br><br>RNA-BM: CCGTCGTC | 0.003<br><br>0.0003 | 2.89<br><br>0.08 | NAC_tnt.AT3G12910_col_a_m1: NAC (No Apical Meristem) domain transcriptional regulator superfamily protein, regulates LHY expression in response to stress[8]. Involved in hormone signalling and biotic/abiotic stress responses. Diurnal oscillation of transcript.<br><br>RNA-binding protein: RNCMPT00073 (SRSF7- Serine And Arginine Rich Splicing Factor 7)[9]<br><br>miRNA targets: miR8175 (chloroplast biogenesis)[10] |
| 4 | TTTCAG.1 | mRNA UTR:40.71% | TTTCAA | 0.003 | 2.13 | ND_tnt.AT1G63040_col_a_m1: pseudogene member of DREB subfamily A-4 of ERF/AP2 transcription factor family. |
| 5 | ATAAAA.1 | mRNA UTR:81.30% | ATAAAACGT | 0.004 | 0.2 | SORLREP2: Overrepresented in PHYA-repressed promoters[11] (light repressed sequences) |
| 6 | TCTCTC.1 | mRNA UTR:78.56% | CTCTCTCTCTCTCTC<br><br>CTCACTC | 0.00004<br><br>0.02 | 0.04<br><br>0.92 | BBRBPC_tnt.BPC1_colamp_a_m1: BASIC PENTACYSTEINE1 (BPC1), belongs to BBR/BPC family. Regulator of the homeotic gene *SEEDSTICK* (*STK*), which controls ovule identity. BioGRID[3]: interacts with BPC6, BPC4, BPC1 and circadian related TCP4, TCP14, TCP7 and TCP15[4]<br><br>SORLIP5 (light-induced sequences) |
| 7 | CCGCCG.1 | mRNA UTR:10.53% | CCACCGCCGCCGCCA | 0.00003 | 0.02 | AP2EREBP_tnt.ERF5_col_a_m1: Ethylene-responsive transcription factor 5 (ERF5), has core link to circadian clock. ChIP-seq revealed the binding of TOC1 and PRR5 proteins to ERF5[12]. Plants constitutively expressing ERF5 display upregulation of pathogen defence genes[13]. Good matches also in ethylene response factors ERF115 (AT5G07310), CRF10 (AT1G68550) and ERF11 (AT1G28370). |

| | | | | | | |
|---|---|---|---|---|---|---|
| | | | TCCTCCG | 0.0006 | 0.155 | RNA-binding protein: RNCMPT00162 (LIN28A)[14] |
| 8 | TCTCTT.1 | mRNA UTR:87.66% | CTCAAGTGA | 0.05 | 2.13 | SORLIP3<br>light-induced sequences |
| 9 | GTGCAA.1 | mRNA UTR:13.44% | TGTGNNATG<br><br>CTTGTGCAACA CGTAA | 0.04<br><br>0.006 | 1.67<br><br>4.83 | CO response element<br><br>NAC_tnt.ANAC058_col_a_m1; NAC DOMAIN CONTAINING PROTEIN 58, DNA-binding transcription factor activity |
| 10 | ATGCAT.1 | mRNA UTR:23.80% | ATATACATGCA TGCA | 0.001 | 0.83 | ABI3VP1_tnt.FUS3_col_a_m1; FUS3, Transcription factor with high similarity to the B3 region of VP1/ABI3-like proteins. FUS3 acts through the RY-promoter element to control gene expression during late embryogenesis and seed development[15] |
| 11 | TATATA.1 | mRNA UTR:68.67% | TGTATATAT | 0.004 | 0.2 | SORLREP3<br>light repressed sequences |
| 12 | AAATAA.1 | mRNA UTR:76.01% | ATAAATAACGG TATT | 0.001 | 1.21 | MYB_tnt.MYB56_col_a_m1: MYB56, negative regulator of flowering[16]. AtMyb56 regulates the expression of AtGPT2 in response to circadian rhythms and sucrose thereby regulating Anthocyanin Levels[17] |
| 13 | TAAAAA.1 | mRNA<br><br>UTR:83.64% | ACGGTAAAAA<br><br>ACGGTAAAAAA<br><br>TAAAAATATCT | 0.0005<br><br>0.0007<br><br>0.003 | 0.42<br><br>0.61<br><br>2.2 | Trihelix_tnt.GTL1_col_a_m1: GTL1, GT2-LIKE 1. Regulates ploidy-dependent cell growth in trichome.<br><br>Trihelix_tnt.GT2_colamp_a_m1: GT2, evening element binding factor[18]<br>MYBrelated_tnt.LHY1_colamp_a_m1: LHY1 circadian clock gene[18] |
| 14 | AAGCTC.1 | mRNA UTR:31.04% | CTTCTAGAAGC TT | 0.011 | 9.64 | HSF_tnt.HSFB3_colamp_a_m1: HSFB3-heat stress transcription factor B-3. Transcriptional regulator that specifically binds heat shock promoter elements (HSE); Belongs to the HSF family. BioGRID[3] Interacts with: ERF4, TCP9[4], TCP4[4], AGL17[19], AGL16[19], TCP10[4], TCP13[4], TCP14[4] |
| 15 | TGGAGA. 1 | mRNA UTR:32.70% | TACCTGGAGAA CATA | 0.001 | 0.82 | C2H2_tnt.WIP5_col_a_m1:<br>WIP5, DNA binding transcription factor activity |
| 16 | ATGGAG. 1 | mRNA UTR:27.21% | GATGATGGA | 0.006 | 5.46 | CCAATHAP3_tnt.HAP3_col_a_m1:<br>At HAP2 or At HAP3 overexpression may impair formation of a CO/At HAP3/At HAP5 complex leading to reduced expression of FT[20] |
| 17 | GGTGTT.1 | mRNA UTR:22.97% | TTGTGCAGGTG TTATCTG | 0.002 | 1.6 | C2H2_tnt.At5g22890_col_a_m1: SENSITIVE TO PROTON RHIZOTOXICITY 2, STOP2. A homolog of AT-STOP1 responds to acidic pH to activate a malate efflux transporter with cycling expression, suggesting regulation of CAM photosynthesis[21] |
| 18 | CTGCTG.1 | mRNA UTR:12.96% | TGCTGCTGCTG CTGC | 0.00009 | 0.08 | NLP_tnt.AtNLP4_col_b_m1: NLP4 NLP Transcription factor, has protein-protein interaction with PP5 (phytochrome-specific type 5 serine/threonine protein phosphatase). Dephosphorylates active Pfr-phytochromes. Controls light signal flux by enhancing phytochrome stability and affinity for a signal transducer. BioGRID[3] Interacts with: AGL15[22]-a repressor of floral transition in Arabidopsis.<br><br>miRNA targets; miR172(2)(developmental timing)[23] |
| 19 | TTCACT.1 | mRNA UTR:43.50% | ACTTCACT<br><br>GAGTGAG | 0.0002<br><br>0.02 | 0.21<br><br>1.17 | C2H2_tnt.STZ_colamp_a_m1; C2H2 zinc finger STZ, involved in JA signalling, response to photooxidative stress and associated with RAV1[24]<br><br>SORLIP5 (light-induced sequences) |
| 20 | CACCGT.1 | mRNA UTR:12.63% | CCTCCACCGTC CAT<br><br>CACCGCG | 0.001<br><br>0.01 | 0.69<br><br>0.7 | AP2EREBP_tnt.At1g19210_col_a_m1; ERF17 a member of the DREB subfamily A-5 of ERF/AP2 transcription factor family<br><br>FHY1-FAR1 binding sites |
| 21 | TCTCTG.1 | mRNA UTR:55.73% | AAACTGATATC TCTGTC | 0.002 | 3.81 | UP00080_2: UP00080_2 is a motif from mice representing GATA1, GATA TRANSCRIPTION FACTOR 1, GO biological process: Circadian |

| | | | | | | |
|---|---|---|---|---|---|---|
| | | | | | | rhythm, regulation of transcription. GATA1 in Arabidopsis is circadian but recognises motif GATAAGG[25] |
| 22 | GAAGAC | Promoter | TGAAAAC | 0.005 | 4.19 | CCAATHAP3_tnt.NFYB4_col_a_m1; NFYB4, Nuclear transcription factor Y subunit B-4, belongs to HAP3/NFYB subunit family-overexpression reduces FT expression[20] |
| 23 | GGTATA.1 | mRNA UTR:14.12% | AAAGTTAGGTATAA | 0.003 | 2.37 | MYB_tnt.MYB121_col_a_m1; MYB121 a putative transcription factor, member of the R2R3 factor gene family. MYB121 homeolog in interacts with RNA recognition motif (RRM)-containing protein; SR-like splicing factor required for phytochrome B (phyB) signal transduction and involved in phyB-dependent alternative splicing |
| | | | TGTATATAT | 0.05 | 2.16 | SORLREP3 (light repressed sequences) |
| 24 | CTCTCT.1 | mRNA UTR:75.73% | CTCTCTCTCTCTCTC | 0.00004 | 0.04 | BBRBPC_tnt.BPC1_colamp_a_m1: BASIC PENTACYSTEINE1 (BPC1) , belongs to BBR/BPC family. Regulator of the homeotic gene *SEEDSTICK* (*STK*), which controls ovule identity. BioGRID[3]: Interacts with: BPC6, BPC4, BPC1 and circadian related TCP4, TCP14, TCP7 and TCP15[4] |
| | | | GAGTGAG | 0.02 | 0.92 | SORLIP5 (light-induced sequences) |
| 25 | CGTCTC.1 | mRNA UTR:22.17% | CGTATC | 0.009 | 0.43 | Antisense LUX Binding Site |
| 26 | TGATTG | Promoter | TCAATGATTGATT | 0.0002 | 0.15 | Homeobox_tnt.ATHB7_col_a_m1; HOMEOBOX 7 (ATHB7) rapidly induced by cold stress[26], it encodes a putative transcription factor that contains a homeodomain closely linked to a leucine zipper motif. Regulated in an ABA-dependent manner and may act in a signal transduction pathway to mediate drought response. |
| | | | AGATTGTT | 0.04 | 1.69 | CCA1 binding site |
| 27 | TGATCA.1 | mRNA UTR:36.74% | - | - | - | - |
| 28 | CGTTTT | Promoter | ACGTTTTAT | 0.002 | 0.106 | SORLREP2 (light repressed sequences) |
| 29 | CTCTTC.1 | mRNA UTR:61.86% | ATTTGGCTTTTCGGA | 0.01 | 9.02 | RWPRK_tnt.NLP7_col_a_m1; Encodes NIN Like Protein 7 (NLP7). Modulates nitrate sensing and metabolism, Enhancing plant growth in N-sufficient conditions[27] |
| 30 | AACTCT | Promoter | GAAATCT | 0.008 | 7.37 | C2H2_tnt.AT3G49930_col_a_m1; C2H2/C2HC zinc fingers superfamily protein |
| | | | AAACCCT | 0.02 | 1.17 | Telo-box (TBX) |

**Table S6. The top 30 most variable features (*k*-mers), by SHAP explanation values, between the Arabidopsis true positive circadian genes classed as morning, day, evening and night phase.** For our best performing classifier LightGBM we split the genes into morning, day, evening and night and investigated which *k*-mers differentiated the groups. We identified the top 30 most variable *k*-mers between the four groups consensus SHAP explanations, these *k*-mers should therefore vary most in their importance between the groups (see Methods). Here, we detail the closest matches of the *k*-mer's to known Arabidopsis transcription factor binding sites with (p<0.05) as defined using Tomtom motif comparison. The position of the *k*-mer is described as promoter or mRNA; if mRNA we express the percentage of mRNAs with the *k*-mer in their UTR region as a proportion of all genes with a UTR k-mer. Features in bold type were not observed in Table S5.

| | K-mer | Position | Associated Motif | Description |
|---|---|---|---|---|
| 1 | TCTCCG.1 | mRNA UTR:28.18% | AAATCTCCGGCGACG | Trihelix_tnt.AT3G58630_col_a_m1 (Table ZB) |
| 2 | TCTTCT.1 | mRNA UTR:93.34% | GGAATCTTCTA | G2like_tnt.At1g25550_col_m1 (Table ZB) |
| 3 | CCGTCG.1 | mRNA UTR:11.23% | CCGTCGAATTCACGGCA | NAC_tnt.AT3G12910_col_a_m1 (Table ZB) |
| 4 | TCTCTC.1 | mRNA UTR:78.56% | CTCTCTCTCTCTCTC | BBRBPC_tnt.BPC1_colamp_a_m1 (Table ZB) |
| 5 | CCGCCG.1 | mRNA UTR:10.53% | CCACCGCCGCCGCCA | AP2EREBP_tnt.ERF5_col_a_m1 (Table ZB) |
| 6 | ATAAAA.1 | mRNA UTR:81.30% | ATAAAACGT | SORLREP2 (light repressed sequences) (Table ZB) |
| 7 | GGTGTT.1 | mRNA UTR:22.97% | TTGTGCAGGTGTTATCTG | C2H2_tnt.At5g22890_col_a_m1 (Table ZB) |
| 8 | ATGGAG.1 | mRNA UTR:27.21% | GATGATGGA | CCAATHAP3_tnt.HAP3_col_a_m1 (Table ZB) |
| **9** | **GATATT** | **Promoter** | **AGATATTTT** | **Evening element (p=0.0016, E=0.074)** |
| **10** | **AAACCC.1** | **mRNA UTR:41.37%** | **AAACCCT** | **Telo-box TBX (p=0.0008, E=0.038)** |
| **11** | **GCTGAG.1** | **mRNA UTR:13.02%** | **GAAATGGCGGCGGAG** | **AP2EREBP_tnt.ERF13_col_b_m1 (p=0.003,E=2.83); ERF13 ethylene-responsive element binding factor 13** |
| **12** | **CCACGT** | **Promoter** | **CCACGT** | **G-box related sequence (p=0.0003, E=0.014)** |
| **13** | **CTGGAG.1** | **mRNA UTR:11.21%** | **TACCTGGAGAACATA** | **C2H2_tnt.WIP5_col_a_m1 (p=0.0006, E=0.49); DNA binding transcription factor activity** |
| 14 | AAATAA.1 | mRNA UTR:76.01% | ATAAATAACGGTATT | MYB_tnt.MYB56_col_a_m1 (Table ZB) |
| **15** | **ACTGAT** | **Promoter** | **ACTGGT** | **bHLH_tnt.bHLH64_col_a_m1 (p=0.0029, E=2.507); AtbHLH64. Direct binding and activation of EXPA1 and EXPA8. Blue and red light upregulate α-expansin 1 (*EXPA1*) in *Brassica rapa* and its overexpression promotes leaf and root growth in Arabidopsis[28]. BioGRID[3] interacts with: ERF10, TCP8, TCP10, TCP23, PAR1, BEE2, TCP14, TCP15[29], MYB56[30] and ARF6** |
| 16 | TCTCTT.1 | mRNA UTR:87.66% | CTCAAGTGA | SORLIP3 (light-induced sequences) (Table ZB) |
| 17 | TGGAGA.1 | mRNA UTR:32.70% | TACCTGGAGAACATA | C2H2_tnt.WIP5_col_a_m1 (Table ZB) |
| 18 | GTGCAA.1 | mRNA UTR:13.44% | TGTGNNATG | CO response element (Table ZB) |
| **19** | **AATATC.1** | **mRNA UTR:39.23%** | **AAAATATCT** | **Evening element (p=0.0016, E=0.073)** |
| **20** | **GCTGTC** | **Promoter** | **CCAGCTGTCAT** | **bZIP_tnt.bZIP18_colamp_a_m1 (p=0.0004, E=0.239); bZIP18 transcription factor involved in plant development, osmosensory signaling and stress response[31]** |
| **21** | **ACCGGT.1** | **mRNA UTR:12.83%** | **ACCAGT** | **bHLH_tnt.bHLH64_col_a_m1 (p=0.0015, E=1.307); Direct binding and activation of EXPA1 and EXPA8. Blue and red light upregulate α-expansin 1 (*EXPA1*) in *Brassica rapa* and its overexpression promotes leaf and root growth in Arabidopsis[28]. BioGRID[3] interacts with: ERF10, TCP8, TCP10, TCP23, PAR1, BEE2, TCP14, TCP15[29], MYB56[30] and ARF6** |

| | | | | |
|---|---|---|---|---|
| **22** | **AGATGA.1** | **mRNA UTR:37.65%** | **GGAAGATGAAACGTC** | **RWPRK_tnt.RKD2_colamp_a_m1 (p=0.0030, E=2.630); RKD2, RWP-RK domain containing protein. Role in egg cell diffeentiation[32]** |
| **23** | **GACCCG.1** | **mRNA UTR:5.49%** | **TGGTGGACCCA** | **TCP_tnt.TCP16_colamp_a_m1 (p=0.0021, E=1.809); TCP16 involved in Leaf and pollen development. BioGRID[3] interacts with: ERF4, ERF10, ERF112, ARF19, AXR3, RVE2[6]** |
| **24** | **AAGAAG.1** | **mRNA UTR:70.99%** | **AAACTTGTAAAAGAAGTAA** | **NAC_tnt.ANAC005_col_a_m1 (p=0.0019, E=1.700); ANAC005, membrane associated transcription factor regulates vascular development[33]** |
| **25** | **TGGGCC** | **Promoter** | **TGTGGGCCCCACTT** | **TCP_tnt.At5g08330_col_a_m1 (p=0.0002, E=0.215); TCP21, Circadian oscillator protein which interacts with bZIP63 and regulates a response of CCA1 to sugars[34]** |
| **26** | **TTCTTC.1** | **mRNA UTR:91.02%** | **GTTTCTTCATCTTCAAGTA** | **NAC_tnt.NTL8_col_m1 (p=0.002, E=1.89); NTL8 mediates salt-responsive flowering via *FT* in Arabidopsis and that membrane-mediated transcription regulation underlies the salt signaling in mediating flowering initiation[35]** |
| 27 | TGATCA.1 | mRNA UTR:36.74% | - | (Table ZB) |
| **28** | **TGGTCT.1** | **mRNA UTR:23.99%** | **GTGGTCCCCAC** | **TCP_tnt.TCP17_colamp_a_m1 (p=0.009, E=8.25); (TCP17). TCP5, TCP13, and TCP17, promote thermoresponsive hypocotyl growth by positively regulating PIF4. TCP17 interacts with a blue light receptor, CRYPTOCHROME 1 (CRY1), at lower temperature, leading to reduced activity of TCP17[36]. BioGRID[3] interacts with: TCP2, TCP4, TCP5, TCP8, TCP10, TCP11, TCP13, TCP14, TCP19, TCP20, TCP21, TCP23, TCP24, ERF107, ERF112, RVE2[6]** |
| 29 | TTCACT.1 | mRNA UTR:43.50% | ACTTCACT | C2H2_tnt.STZ_colamp_a_m1 (Table ZB) |
| **30** | **CTCGCC.1** | **mRNA UTR:8.93%** | **CGTCGCCGGAGATTT** | **Trihelix_tnt.AT3G58630_col_a_m1 (p=0.0065, E=5.69); AT3G58630 has a protein-protein interaction[1] with LIGHT INSENSITIVE PERIOD1, LIP1 which influences the light input pathway of the plant circadian network[2]. Interacts[2] with RIN1-a component of the phosphoprotein phosphatase 2A regulatory subunit A. Phosphorylation of circadian clock proteins is an essential posttranscriptional mechanism in their regulation.** |

**Table S7. Comparing the average SHAP value for *k*-mers of interest between genes defined as morning, day evening and night phase from the Arabidopsis true positive circadian genes.** For our best performing classifier LightGBM we split the genes into morning, day, evening and night and investigated average *k*-mer SHAP values for each of the groups. Here, we detail t tests to compare the groups for *k*-mers defined as having a high variation between morning, day, evening and night groups.

| Motif/Element | *K*-mer Sequence | Comparison 1 | Mean SHAP value | Comparison 2 | Mean SHAP value | t value | p-value | Significantly different |
|---|---|---|---|---|---|---|---|---|
| EE | GATATT | Morning | 0.0098 | Evening | 0.0220 | 2.9882 | 0.0029 | Yes |
| EE | GATATT | Day | 0.0180 | Evening | 0.0220 | 1.0272 | 0.3046 | No |
| EE | GATATT | Night | 0.0088 | Evening | 0.0220 | 3.5706 | 0.0004 | Yes |
| EE | AATATC.1 | Morning | 0.0053 | Evening | 0.0149 | 2.1517 | 0.0318 | Yes |
| EE | AATATC.1 | Day | 0.0100 | Evening | 0.0149 | 1.1280 | 0.2596 | No |
| EE | AATATC.1 | Night | 0.0019 | Evening | 0.0149 | 3.8316 | 0.0001 | Yes |
| Telo-box | AAACCC.1 | Morning | 0.0116 | Night | 0.0176 | 1.4020 | 0.1613 | No |
| Telo-box | AAACCC.1 | Day | 0.0030 | Night | 0.0176 | 4.6346 | 0.0001 | Yes |
| Telo-box | AAACCC.1 | Evening | 0.0063 | Night | 0.0176 | 2.6060 | 0.0093 | Yes |
| G-box | CCACGT | Day | 0.0084 | Morning | 0.0126 | 0.7381 | 0.4606 | No |
| G-box | CCACGT | Evening | 0.0182 | Morning | 0.0126 | 0.7366 | 0.4617 | No |
| G-box | CCACGT | Night | 0.0038 | Morning | 0.0126 | 1.6908 | 0.0912 | No |

**Table S8. Comparing accuracy for MetaCycle, our ML model using 12 timepoints and our ML model using DNA sequence, considering PHYA-E.** Accuracies reported are all in the format number of correctly classified genes/total number of genes classified.

| Name | PHYA | PHYB | PHYC | PHYD | PHYE | Accuracy for PHYA-E |
|---|---|---|---|---|---|---|
| Transcript | AT1G09570.1 | AT2G18790.1 | AT5G35840.1 | AT4G16250.1 | AT4G18130.1 | - |
| Literature | Circadian | Circadian | Low amplitude-potentially Circadian | Circadian | Circadian | 100% |
| MetaCycle | Circadian | Circadian | Non-Circadian | Non-Circadian | Non-Circadian | 40% |
| ML best model with 12TP | Circadian | Circadian | Non-circadian | Circadian | Circadian | 80% |
| K-mer best model | Circadian | Circadian | Non-circadian | Circadian | Circadian | 80% |
| Were genes used in ML model training | Yes | Yes | No | No | No | - |

**Table S9. Results from SHAP explanation of Arabidopsis (Col-0) circadian/non-circadian best ML binary classification DNA sequence-based analysis.** Showing the top three computed values when we compared the SHAP explanations between each of the PHY genes A/B/D/E and PHYC i.e. PHYA versus PHYC, PHYB versus PHYC, PHYD versus PHYC and PHYD versus PHYC.

| | **PHYA-C** | | **PHYB-C** | | **PHYD-C** | | **PHYE-C** | |
|---|---|---|---|---|---|---|---|---|
| Rank | K-mer | SHAP difference | K-mer | SHAP difference | K-mer | SHAP difference | K-mer | SHAP difference |
| 1 | GGTAGA.1 | 0.458342219 | TCTCCG.1 | 0.80413333 | CCGTCG.1 | 0.49510482 | GGTAGA.1 | 0.45366824 |
| 2 | TTTCTG.1 | 0.211106645 | GGTAGA.1 | 0.46381106 | TTTCAG.1 | 0.2964691 | TCTCCG.1 | 0.4508364 |
| 3 | AAATAA.1 | 0.195492878 | CCGTCG.1 | 0.3521632 | GGTAGA.1 | 0.28438412 | TTTCTG.1 | 0.20130146 |

**Table S10. List of 41 well-known circadian genes**. Known circadian genes that have evidence for rhythmicity either from published reports or where that is not clear, experimental evidence from the Romanowski et al. (2020) dataset is used to confirm (MetaCycle q<0.05).

| Gene | Description | Gene | Description |
|---|---|---|---|
| AT2G25930 | ELF3 | AT3G09600 | RVE8 |
| AT2G40080 | ELF4 | AT5G15840 | CO |
| AT3G46640 | LUX | AT1G04400 | CRY2 |
| AT5G59570 | BOA | AT2G32950 | COP1 |
| AT1G68050 | FKF1 | AT1G12910 | LWD1 |
| AT4G08920 | CRY1 | AT3G26640 | LWD2 |
| AT5G61380 | TOC1 | AT1G29930 | CAB1 |
| AT3G22380 | TIC | AT1G29910 | CAB2 |
| AT5G08330 | CHE | AT4G33980 | COR28 |
| AT1G22770 | GI | AT5G42900 | COR27 |
| AT1G01060 | LHY | AT3G54500 | LNK2 |
| AT2G46830 | CCA1 | AT5G64170 | LNK1 |
| AT5G60100 | PRR3 | AT3G13550 | COP10 |
| AT5G24470 | PRR5 | AT4G10180 | DET1 |
| AT5G02810 | PRR7 | AT4G16250 | PHYD |
| AT2G46790 | PRR9 | AT4G18130 | PHYE |
| AT2G43010 | PIF4 | AT4G31120 | PRMT5 |
| AT2G20180 | PIF1 | AT1G67840 | CSK |
| AT1G09570 | PHYA | AT3G20810 | JMJ30 |
| AT2G18790 | PHYB | AT5G24120 | SIG5 |
| AT5G02840 | RVE4 | | |

**Table S11. Comparing accuracy for MetaCycle, our ML model using 12 timepoints and our ML model using DNA sequence, considering the 41 well-known circadian genes.** Accuracies reported are all in the format number of correctly classified genes/total number of genes classified.

| Test | Accuracy for PHYA-E | Overall accuracy for 41 circadian | Overall accuracy for 10 circadian (not in train) |
|---|---|---|---|
| Were genes used in Col-0 ML model training? | Yes (40.00%) | Yes (75.61%) | No |
| Evidence (assumed 100%) | 100% | 100% | 100% |
| Col-0<br>MetaCycle | 40% | 80.49% | 20.00% |
| Col-0<br>ML best model with 12TP | 80% | 95.12% | 80.00% |
| Ws-2<br>ML best model with 2TP | 100% | 72.50% | 50.00% |
| Col-0<br>K-mer best model | 80% | 92.68% | 90.00% |
| Ws-2<br>K-mer best model | 60% | 82.50% | 66.66% |
| Ws-2<br>K-mer best model (only labels passing 0.8 threshold considered) | 100% (2 genes) | 90.32% (31 genes) | 83.33% (6 genes) |

**Table S12. Transcripts identified as circadian for Col-0 and non-circadian for Ws-2 by the DNA sequence-based model.** These transcripts are ranked in descending order according to likelihood (increasing probability of circadian in Col-0 versus decreasing probability of non-circadian in Ws-2). Col-0 is classified circadian and Ws-2 is classified non-circadian meeting the following criteria; from the DNA sequence-based model both classifications have a predictive probability >0.8, from the Ws-2 transcriptomic data the same classification is observed with a predictive probability of >0.8 (2 timepoints Graf et al. 2017) and finally, also from the Col-0 transcriptomic data (12 timepoints Romanowski et al., 2020 and 2 timepoints Graf et al. 2017) the same classification is observed.

| Transcript | Description |
|---|---|
| AT1G58602.1 | RECOGNITION OF PERONOSPORA PARASITICA 7, RPP7 |
| AT4G16990.15 | RESISTANCE TO LEPTOSPHAERIA MACULANS 3, RLM3 |
| AT1G58200.1 | MSCS-LIKE 3, MSL3<br>A member of MscS-like gene family, structurally very similar to MSL2, comprising of an N-terminal chloroplast transit peptide, five trans-membrane helices and a C-terminal cytoplasmic domain. Mutant plants showed abnormalities in the size and shape of plastids. MSL3-GFP was localized to discrete foci on the plastid envelope and co-localize with the plastid division protein AtMinE. MSL3 was capable of increasing the osmotic-shock survival of a mutant bacterial strain lacking MS-ion-channel activity. |
| AT4G23250.2 | CRK17, CYSTEINE-RICH RLK (RECEPTOR-LIKE PROTEIN KINASE) 17, DUF26-21, EMB1290, EMBRYO DEFECTIVE 1290, RECEPTOR-LIKE KINASE THALIANACOL-0GENOMICLIBRARY(CLONTECH).THEPOSITIVEPHAGECLONES C-X8-C-X2-C CLASS 1, RKC1 |
| AT5G35180.1 | ENHANCED DISEASE RESISTANCE protein (DUF1336); |
| AT1G17460.1 | TRF-LIKE 3, TRFL3: Arabidopsis thaliana myb family transcription factor (At1g17460) |
| AT3G57410.9 | ATVLN3, VILLIN 3, VLN3; Encodes a protein with high homology to animal villin. VLN3 is a Ca2+-regulated villin involved in actin filament bundling. |
| AT1G09300.1 | ATICP55; Encodes a mitochondrial protease ICP55. Alters the stability of proteins by removal of a single amino acid from their sequence. |
| AT1G24706.7/.3 | ATTHO2, EMB2793, EMBRYO DEFECTIVE 2793, THO2; Encodes a component of the putative Arabidopsis THO/TREX complex: THO1 or HPR1 (At5g09860), THO2 (At1g24706), THO3 or TEX1 (At5g56130), THO5 (At5g42920, At1g45233), THO6 (At2g19430), and THO7 (At5g16790, At3g02950). THO/TREX complexes in animals have been implicated in the transport of mRNA precursors. Mutants of THO3/TEX1, THO1, THO6 accumulate reduced amount of small interfering (si)RNA, suggesting a role of the putative Arabidopsis THO/TREX in siRNA biosynthesis. Mutations in THO have severe developmental defects and affect the production of several different classes of small RNAs indicating a broader role in small RNA biosynthesis. |
| AT5G27360.1 | SFP2; Encodes a sugar-porter family protein that unlike the closely related gene, SFP1, is not induced during leaf senescence |
| AT3G52050.6 | OEX1, ORGANELLAR EXONUCLEASE 1; 5-3 exonuclease family protein |

**Table S13. Using transcriptomic information to predict circadian time.** Mean absolute errors across the three datasets for different numbers of genes before hyperparameter optimisation

| Error (MAE) | 1000 genes | 100 genes | 50 genes | 10 genes | 5 genes |
|---|---|---|---|---|---|
| Training | 9 minutes | 4minutes | 44 minutes | 11 minutes | 10 minutes |
| Validation | 125 minutes | 81 minutes | 119 minutes | 84 minutes | 125 minutes |
| Test | 180 minutes | 289 minutes | 104 minutes | 146 minutes | 125 minutes |

**Table S14. List of 15 selected genes for prediction of circadian time.**

| Gene | Description |
|---|---|
| AT1G13650.1 | |
| AT3G55450.1 | PBL1 |
| AT1G02930.2 | GSTF6 |
| AT1G79500.3 | AtkdsA1 |
| AT5G24850.1 | CRY3 |
| AT5G06870.1 | PGIP2 |
| AT5G01820.1 | SR1 |
| AT4G08870.1 | ARGAH2 |
| AT1G75100.1 | JAC1 |
| AT2G29650.2 | PHT4 |
| AT5G06690.1 | WCRKC1 |
| AT3G17609.2 | HYH |
| AT4G15690.1 | GRXS5 |
| AT5G41460.1 | |
| AT1G06040.1 | STO |

**Table S15. Parameter training during hyperparameter optimization for comparison of commonly used deterministic ML methods.** Showing the parameters that were tuned using Grid Search and the range of trialled hyperparameters.

| REGRESSOR | HYPERPARAMETER TUNING |
|---|---|
| LOGISTIC REGRESSION | penalty:['l1', 'l2']<br>C:[1.0, 0.5, 0.1]<br>solver:['liblinear'] |
| RANDOM FOREST | criterion: ['gini', 'entropy']<br>min_samples_leaf: [1, 2, 3, 4, 5, 6, 7, 8, 9, 10]<br>n_estimators:[ int(x) for x in np.linspace(start = 100, stop = 4000, num = 10)]<br>max_depth: [1, 2, 3, 4, 5, 6, 7, 8, 9, 10]<br>min_samples_split: [2, 5, 10], max_features:['auto', 'sqrt'] |
| SVM | kernel: ['linear', 'rbf']<br>gamma: scipy.stats.expon(scale=.1)<br>C: scipy.stats.expon(scale=100) |
| KNN | n_neighbors: [1, 2, 3, 4, 5, 6, 7, 8, 9, 10, 11, 12, 13, 14, 15, 16, 17, 18, 19, 20]<br>leaf_size: [1, 2, 3, 4, 5]<br>weights:['uniform', 'distance']<br>algorithm:['auto', 'ball_tree','kd_tree','brute'] |
| XGBOOST | max_depth: [2, 3, 4, 5, 6, 7, 8, 9, 10, 15, 20]<br>subsample: [0.2, 0.5, 0.6, 0.7, 0.8, 0.9, 1]<br>learning_rate: scipy.stats.expon(scale=1)<br>min_child_weight: scipy.stats.expon(scale=10)<br>max_delta_step: [0, 1, 2] |
| DECISION TREE | criterion: ['gini', 'entropy']<br>min_samples_leaf: [1, 2, 3, 4, 5]<br>max_depth: [1, 2, 3, 4, 5]<br>min_samples_split: [2, 3, 4, 5]<br>presort: [True,False] |
| LIGHT GBM | num_leaves: [10, 20, 50, 100, 200]<br>subsample: [0.2, 0.5, 0.6, 0.7, 0.8, 0.9, 1]<br>min_data_in_leaf: [10, 25, 50, 75, 100]<br>max_depth: [3, 5, 6, 7, 8, 10, 15, 20, 25]<br>learning_rate: scipy.stats.expon(scale=1)<br>n_estimators: [50, 150, 200, 250, 300, 350, 400, 450, 500] |
| GAUSSIAN PROCESS | normalize_y:[True,False]<br>copy_X_train:[True, False]<br>alpha: [1e-2, 1e-4, 1e-6, 1e-8, 1e-10, 1e-12]<br>n_restarts_optimizer: [10]<br>WhiteKernel(noise_level=1e-7)<br>length_scale: np.ones(X_train.shape[1]) |
